# Development of Auditory and Spontaneous Movement Responses to Music over the First Postnatal Year

**DOI:** 10.1101/2025.05.04.649695

**Authors:** Trinh Nguyen, Félix Bigand, Susanne Reisner, Atesh Koul, Roberta Bianco, Gabriela Markova, Stefanie Hoehl, Giacomo Novembre

**Affiliations:** Neuroscience of Perception and Action Lab, Italian Institute of Technology, Rome, Italy; Department of Developmental and Educational Psychology, University of Vienna, Vienna, Austria; Department of Developmental and Biological Psychology, Heidelberg University, Heidelberg, Germany; Doctoral School Cognition, Behavior and Neuroscience, University of Vienna, Austria; Department of Translational Research & New Technologies in Medicine and Surgery, University of Pisa, Pisa, Italy; Institute for Early Life Care, Paracelsus Medical University, Salzburg, Austria

**Keywords:** Music, EEG, movement, development, infancy, pitch

## Abstract

Humans across cultures not only share the ability to recognise music but also respond to it through movement. While the sensory encoding of music is well-studied, when and how infants naturally start moving to music is largely unexplored. This study simultaneously investigates infants’ neural (auditory) responses and spontaneous movements to music during the first postnatal year. Neural activity (EEG) and body kinematics (markerless pose estimation) were recorded from 79 infants (aged 3, 6, and 12 months) listening to refrains of children’s music, along with shuffled, high-pitched, and low-pitched versions of the same songs. Neural data revealed that, across all ages, infants exhibit enhanced auditory responses to music compared to shuffled music, indicating that auditory encoding of music emerges early in development. Movement data revealed a different outcome. While coarse auditory-motor coupling is present at all ages, more complex structured movement patterns emerge in response to music only by 12 months. Notably, no age group demonstrated evidence of coordinated movements to music. Additionally, enhanced auditory responses to high vs low pitch were only evident at 6 months, while infants’ movements were better predicted by high-pitched compared to low-pitched music at all ages. This study provides initial insights into how the developing brain gradually transforms music into spontaneous movements of increasing complexity.

## Introduction

Musicality – the biological predisposition to perceive, appreciate, and produce music (Honing, 2018; Trehub, 2003) – is increasingly recognised as a fundamental aspect of human nature. Numerous accounts suggest that engaging with music through movement is at the core of musicality (Honing et al., 2015; Schachner et al., 2009; Trehub et al., 2015). Functionally, such engagement can be broken down into two fundamental components of neurocognitive development: the ability to perceive and recognise music (*sensory component*), and the ability to produce movement responses that are temporally aligned with the musical structure, from coordinated vocalizations and percussive actions up to complex dance moves (*motor component*; Brown, 2022; Trehub, 2003a; Trevarthen, 1999). Despite this inherent predisposition toward music, the developmental trajectory of infants’ musicality remains largely unknown (see Nguyen et al., 2023, for a review). While there is increasing research on infant music perception, including controlled manipulations of select musical features, we know less about the translation of perception into action, namely the ontogenesis of infants’ spontaneous movements to music (see Fujii et al., 2014; Nguyen, Reisner, et al., 2023; Zentner & Eerola, 2010). Furthermore, making our understanding of music-driven motor engagement even more incomplete, no studies to date have looked at both brain activity and spontaneous body movements simultaneously, especially during the first year of life. Accordingly, how the processing of music and its features is transformed into organised motor responses remains underexplored.

The *sensory component* of musicality, namely music perception, can be measured using electroencephalography (EEG), specifically by recording cortical auditory evoked potentials (event-related potentials [ERP]). One of these responses is the infantile P1, a phase-locked EEG positivity peaking around 200-300 ms after an auditory stimulus (Chen et al., 2016; Kushnerenko et al., 2002; Wunderlich et al., 2006). The infantile P1 has been observed in response to both musical notes and speech segments. Auditory evoked potentials, when elicited isochronously, can also be captured using frequency domain analyses (Damsma et al., 2024; Novembre & Iannetti, 2018) such as auditory steady-state responses (ASSR), which are also called steady-state evoked potentials (SSEP, e.g., Cirelli et al., 2016; Nave et al., 2022). These neural responses can provide insight into the developing auditory system and its ability to encode musical structure. Using these neurophysiological measures, prior research has shown that newborns and infants are sensitive to beat structure, pitch deviants and tone interval regularities (Bianco et al., 2025; Edalati et al., 2023; Háden et al., 2009, 2015, 2022; Stefanics et al., 2009; Winkler et al., 2009). Despite these promising results, the neurophysiology of early music processing – particularly its developmental trajectory – remains not fully understood. Here, our primary goal is to investigate infants’ neural encoding of music utilizing both ERP and ASSR approaches to characterize how such neural responses change across the first year of life.

Another component of musicality is the capacity to move to music (*motor component;* Brown, 2022; Fitch, 2015; Honing et al., 2015; Trehub et al., 2015). This capacity is linked to infants not only recognising musical structure but also moving their bodies in response to it. Even though this capacity appears to develop precociously, as evidenced by the fact that even 28-35-week-old foetuses move to music (Kisilevsky et al., 2004), very few studies have systematically examined music-driven spontaneous body movements in infants. An influential paper by Zentner and Eerola (2010) reported that infants across a large age range (from 5 to 24 months) showed more spontaneous rhythmic movements in response to classical music and children’s music compared to infant-directed speech. Importantly, their movements were not synchronized with the musical input, even though a small degree of tempo flexibility was observed (i.e., faster musical tempi evoked relatively faster movement periodicities). The lack of synchrony between music and body movements has also been reported in younger (i.e., 3-4 months old) infants listening to popular music (Fujii et al., 2014). Further, another study testing 7-month-old infants reported more movement in response to (sung) playsongs compared to lullabies but did not assess movement synchrony (Nguyen, Reisner, et al., 2023). Despite these initial investigations, it remains unclear when infants begin to move in response to music, which specific movements are evoked, and when these movements become coordinated with the music. Moreover, a critical limitation in existing research is the lack of a control condition to determine whether these movements are driven specifically by musical structure or reflect general motor activity in response to auditory input. As a second goal, this study is the first to systematically test the gradual development of music-induced movements in different age groups across the first year of life.

Music engages both sensory and motor systems, yet different musical features may differentially shape infants’ engagement with music. While rhythm has been widely studied in early music cognition, pitch is another salient acoustic cue that could play a role in auditory-motor engagement, particularly in infancy. High pitch is a defining feature of infant-directed speech (Fernald & Simon, 1984), among other features such as exaggerated intonation, slower tempo, and simplified vocabulary (Fernald & Kuhl, 1987; Kuhl & Meltzoff, 1982). Similarly, infants most frequently listen to music characterized by high pitch (Costa-Giomi & Sun, 2016; Nakata & Trehub, 2011). Reflecting its prominence, high pitch is found to be one of the most prominent features thought to effectively capture (Conrad et al., 2011; Eckerdal & Merker, 2009; Trainor, 1996; Trainor & Zacharias, 1998) and guide infants’ attention (Lense et al., 2022; Trainor & Desjardins, 2002). On the neural level, infants are also better at encoding pitch deviances in the high voice of polyphonic music, thus showing *high voice superiority* from 3 months of age (Marie & Trainor, 2013, 2014). Taken together, these findings indicate that higher-pitch music would amplify infants’ neural responses (i.e., sensory component) in comparison to lower-pitch music. On the other hand, we know that adults move more to music with greater energy in lower frequencies (Cameron et al., 2022; Stupacher et al., 2013, 2016; Van Dyck et al., 2013). Yet, it remains unknown whether low-pitch music elicits increased movement in infants, as it does in adults, or whether infants’ attraction to high pitch also extends to enhance their motor responses. As a third goal, we thus investigate how musical pitch affects infants’ sensory and motor components.

We presented infants, aged 3, 6, and 12 months, with instrumental refrains of children’s songs (music), shuffled versions of the same songs (shuffled music), and transpositions of the songs that would either emphasize the melody (high pitch) or the bassline (low pitch). We recorded infants’ neural activity using EEG and specifically extracted ERPs and ASSR as indices of infants’ neural response to the various auditory stimuli. We also analysed spontaneous (full-body) movement kinematics using automated video-based motion tracking (DeepLabCut) and extracted *principal movements* using principal component analysis (see Fig. 1, c.f. <u>Bigand et al., 2024</u>, Toiviainen et al., 2010). By adopting a cross-sectional design, we aimed to characterize the maturation of both auditory and movement responses across infancy. We hypothesized that auditory responses would be enhanced when triggered by music compared to shuffled music. This hypothesis was based on the notion that musical structure, notably eroded in the shuffled musical stimuli, is essential to attract infants’ attention towards predictable events (Kouider et al., 2015; Lense et al., 2022). Similarly, based on previous evidence comparing movement responses to music vs speech and silence (Fujii et al., 2014; Zentner & Eerola, 2010), we expected the presence of musical structure to increase the likelihood of spontaneous movements in response to music compared to shuffled music, but we did not have a specific hypothesis about which particular movements would be produced. We further hypothesized that infants would show enhanced neural responses to high-compared to low-pitch music and explored co-occurring differences in spontaneous movements. Generally, we aimed to characterize the maturation of both auditory and motor responses as infants get older. By studying both sensory and motor components of musicality, we aimed to deepen our understanding of when and how infants learn to transform what they perceive into spontaneous movements, eventually leading to the emergence of synchronization to music (Brown, 2022; Fitch, 2015; Honing et al., 2015; Patel & Iversen, 2014).

**Figure 1.**
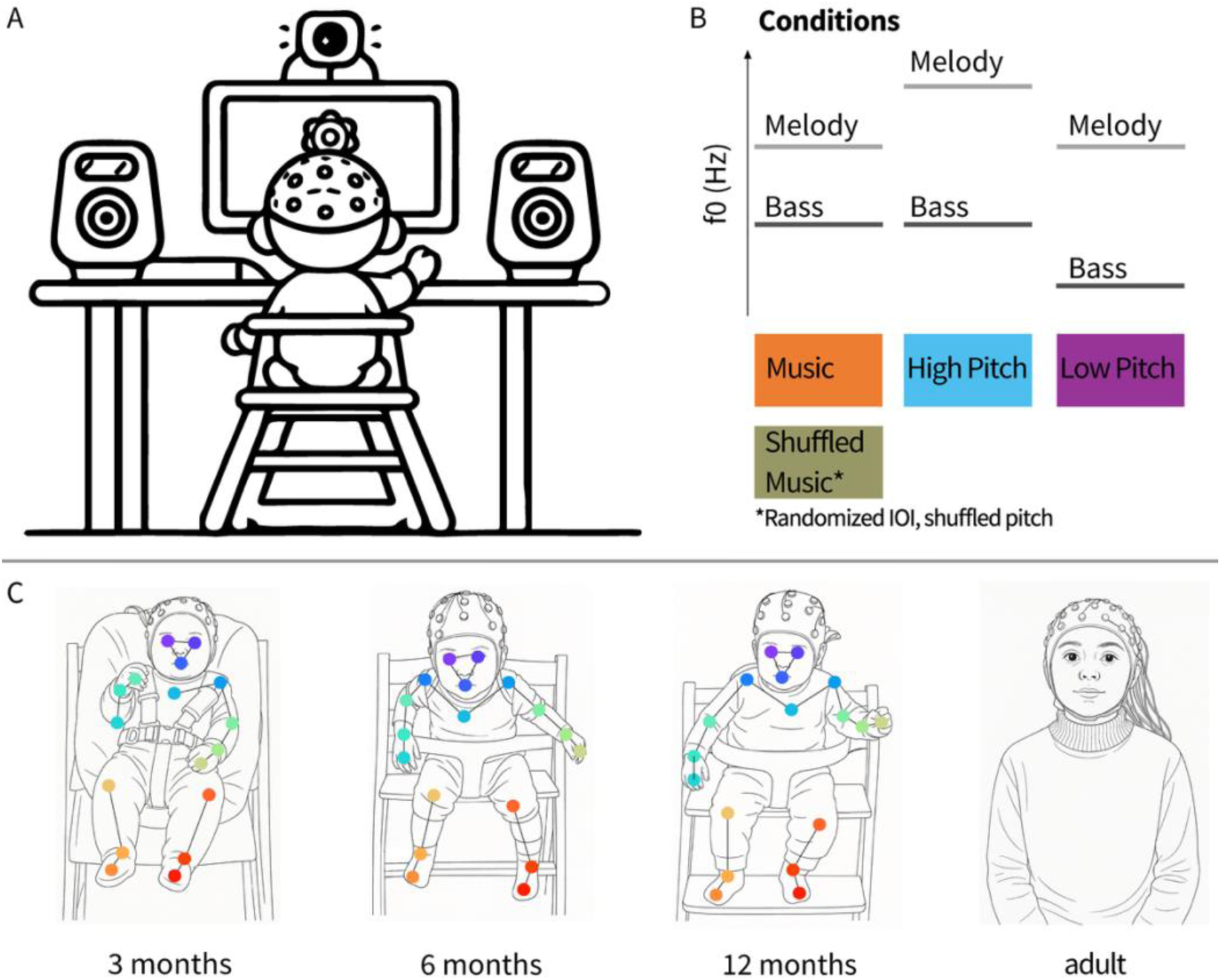
Overview of the procedure (A), experimental conditions (B), and participant sample (C). (A) Infants sat in front of a screen with speakers on each side. The screen showed slowly blossoming flowers to attract infants’ attention. Caregivers (not shown) sat behind the infants and wore noise-cancelling headphones. (B) Infants listened to polyphonic auditory stimuli consisting of a melody and a bassline in four different conditions. The music condition included two children’s songs. The shuffled music condition included versions of the songs used in the music condition that were shuffled in pitch and randomized in inter-onset intervals (IOI). Stimuli belonging to the music and shuffled music conditions had the same pitches. In the high-pitch condition, the melody was shifted one octave higher than in the music condition. In the low-pitch condition, the bassline was shifted one octave lower than in the music condition. Hence, the two voices composing the high-pitch condition were one octave higher than those composing the low-pitch condition. (C) The sample included infants at 3 months (N=26), 6 months (N=26), 12 months (N=27), and an adult control sample (N=26). The dots overlaying the images represent the body parts whose movements were tracked using video-based kinematic analysis.

## Results

This study included EEG and movement measurements taken from 79 infants in the first postnatal year, as well as EEG measurements taken from a control sample of 26 adults. Participants were exposed to two polyphonic children’s songs featuring a melody and a bassline, and their manipulated versions (see Table 1 and S1 for an acoustic characterization of the musical stimuli). We investigated neural responses and movement responses to music vs shuffled music (a control condition in which we shuffled the melody and randomized the inter-onset intervals (IOI) of the music). Additionally, we contrasted neural and movement responses to high- vs low-pitch music. Music vs shuffle conditions (manipulation of structure but not pitch height) and high pitch vs low pitch conditions (manipulation of pitch height but not structure) are contrasted separately to avoid comparing conditions differing in more than one variable. Below, we begin by reporting the neural results, followed by the movement results.

**Table 1.**
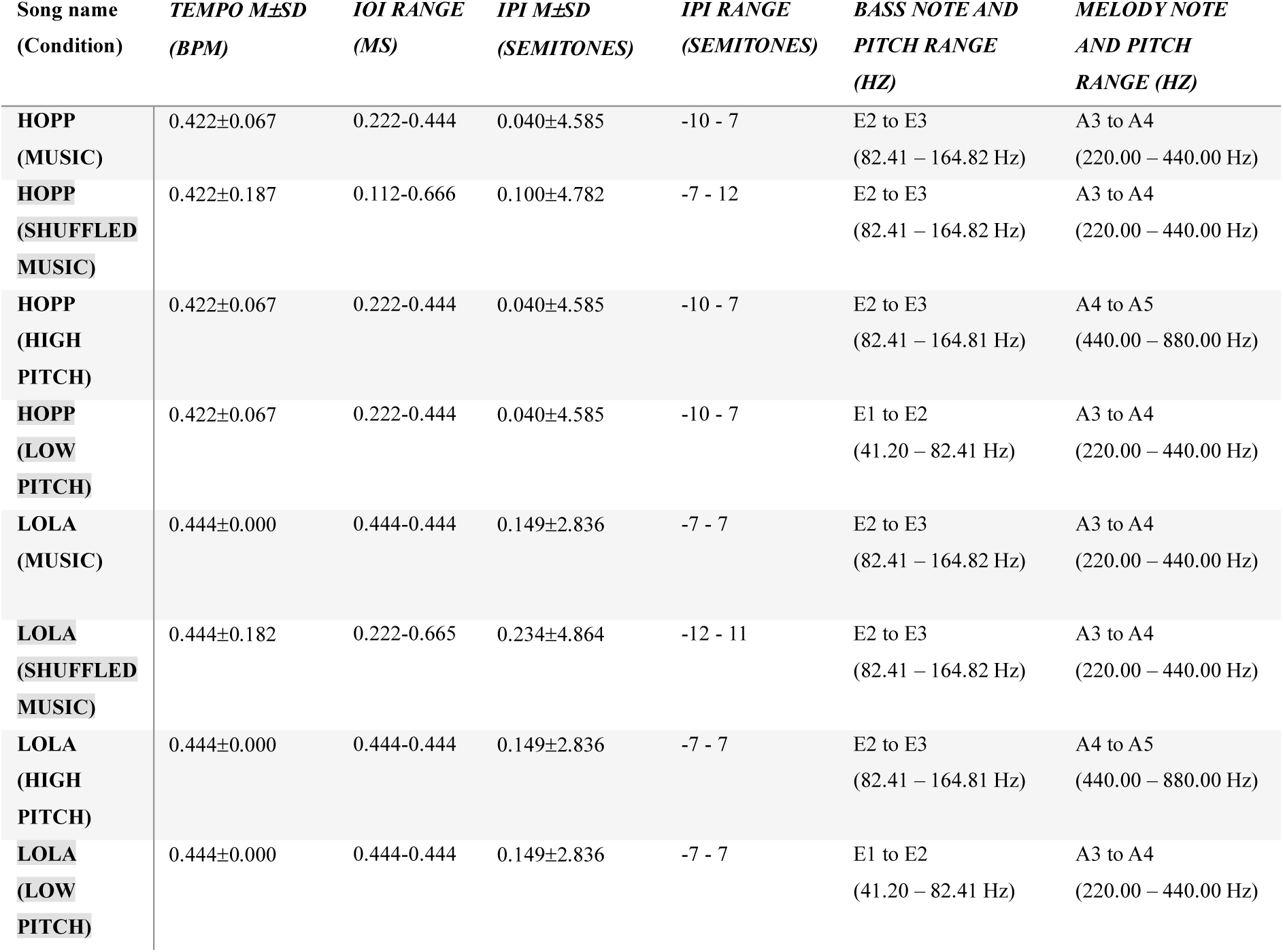
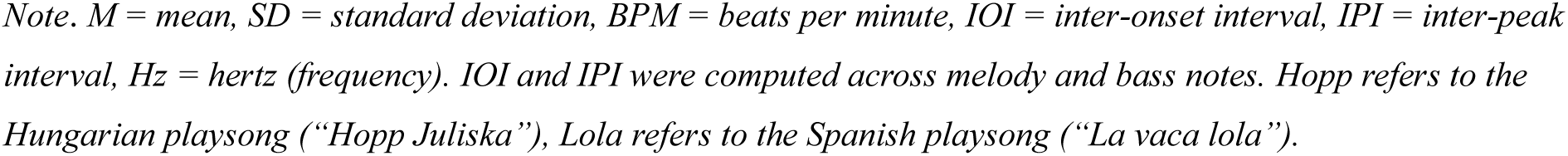
Acoustic characteristics of the musical stimuli.

| Song name<br>(Condition) | TEMPO $M \pm SD$<br>(BPM) | IOI RANGE<br>(MS) | IPI $M \pm SD$<br>(SEMITONES) | IPI RANGE<br>(SEMITONES) | BASS NOTE AND<br>PITCH RANGE<br>(HZ) | MELODY NOTE<br>AND PITCH<br>RANGE (HZ) |
| --- | --- | --- | --- | --- | --- | --- |
| HOPP<br>(MUSIC) | 0.422±0.067 | 0.222-0.444 | 0.040±4.585 | -10 - 7 | E2 to E3<br>(82.41 – 164.82 Hz) | A3 to A4<br>(220.00 – 440.00 Hz) |
| HOPP<br>(SHUFFLED<br>MUSIC) | 0.422±0.187 | 0.112-0.666 | 0.100±4.782 | -7 - 12 | E2 to E3<br>(82.41 – 164.82 Hz) | A3 to A4<br>(220.00 – 440.00 Hz) |
| HOPP<br>(HIGH<br>PITCH) | 0.422±0.067 | 0.222-0.444 | 0.040±4.585 | -10 - 7 | E2 to E3<br>(82.41 – 164.81 Hz) | A4 to A5<br>(440.00 – 880.00 Hz) |
| HOPP<br>(LOW<br>PITCH) | 0.422±0.067 | 0.222-0.444 | 0.040±4.585 | -10 - 7 | E1 to E2<br>(41.20 – 82.41 Hz) | A3 to A4<br>(220.00 – 440.00 Hz) |
| LOLA<br>(MUSIC) | 0.444±0.000 | 0.444-0.444 | 0.149±2.836 | -7 - 7 | E2 to E3<br>(82.41 – 164.82 Hz) | A3 to A4<br>(220.00 – 440.00 Hz) |
| LOLA<br>(SHUFFLED<br>MUSIC) | 0.444±0.182 | 0.222-0.665 | 0.234±4.864 | -12 - 11 | E2 to E3<br>(82.41 – 164.82 Hz) | A3 to A4<br>(220.00 – 440.00 Hz) |
| LOLA<br>(HIGH<br>PITCH) | 0.444±0.000 | 0.444-0.444 | 0.149±2.836 | -7 - 7 | E2 to E3<br>(82.41 – 164.81 Hz) | A4 to A5<br>(440.00 – 880.00 Hz) |
| LOLA<br>(LOW<br>PITCH) | 0.444±0.000 | 0.444-0.444 | 0.149±2.836 | -7 - 7 | E1 to E2<br>(41.20 – 82.41 Hz) | A3 to A4<br>(220.00 – 440.00 Hz) |

*Note. M = mean, SD = standard deviation, BPM = beats per minute, IOI = inter-onset interval, IPI = inter-peak interval, Hz = hertz (frequency). IOI and IPI were computed across melody and bass notes. Hopp refers to the Hungarian playsong (“Hopp Juliska”), Lola refers to the Spanish playsong (“La vaca lola”).*

### EEG: Event-related potentials (ERP)

To examine the neural responses, we analysed the average ERPs time-locked to the acoustic tone onsets of the bassline notes of the auditory stimuli (*Fig. 2*). Adults’ ERPs, which served as a benchmark to interpret infants’ responses, included an early positivity peaking at 37 ms post-stimulus (so-called “P50”, here reaching an average amplitude of 1.05 µV), followed by a later negativity peaking at 87 ms post-stimulus (so-called “N100”, here reaching an amplitude of −0.43 µV) and a second positivity peaking at 158 ms post-stimulus (so-called “P200”, here reaching an amplitude of 0.85 µV). This triphasic EEG pattern has been widely observed in adults in response to fast-rising auditory stimuli and across different contexts (Novembre et al., 2018; Pratt et al., 2008; Remijn et al., 2014; Somervail et al., 2021). Cluster-based permutation analyses, contrasting the ERPs elicited by music vs shuffled music, revealed that the amplitude of the P50 – hereafter referred to as P1 – was larger in response to music compared to shuffled music, particularly within the time range comprised between −17 and 58 ms post-stimulus (*cluster-t*=366.16, *p*=.016). Similarly, the amplitude of the following P200 - hereafter referred to as P2 – was larger in response to music than to shuffled music, particularly within the time range of 114 to 190 ms post-stimulus (*cluster-t*=395.42, *p*=.016). Both P1 and P2 responses were observed over fronto-central electrodes, showing a medial distribution, in line with previous literature (e.g., Lijffijt et al., 2009).

**Figure 2.**
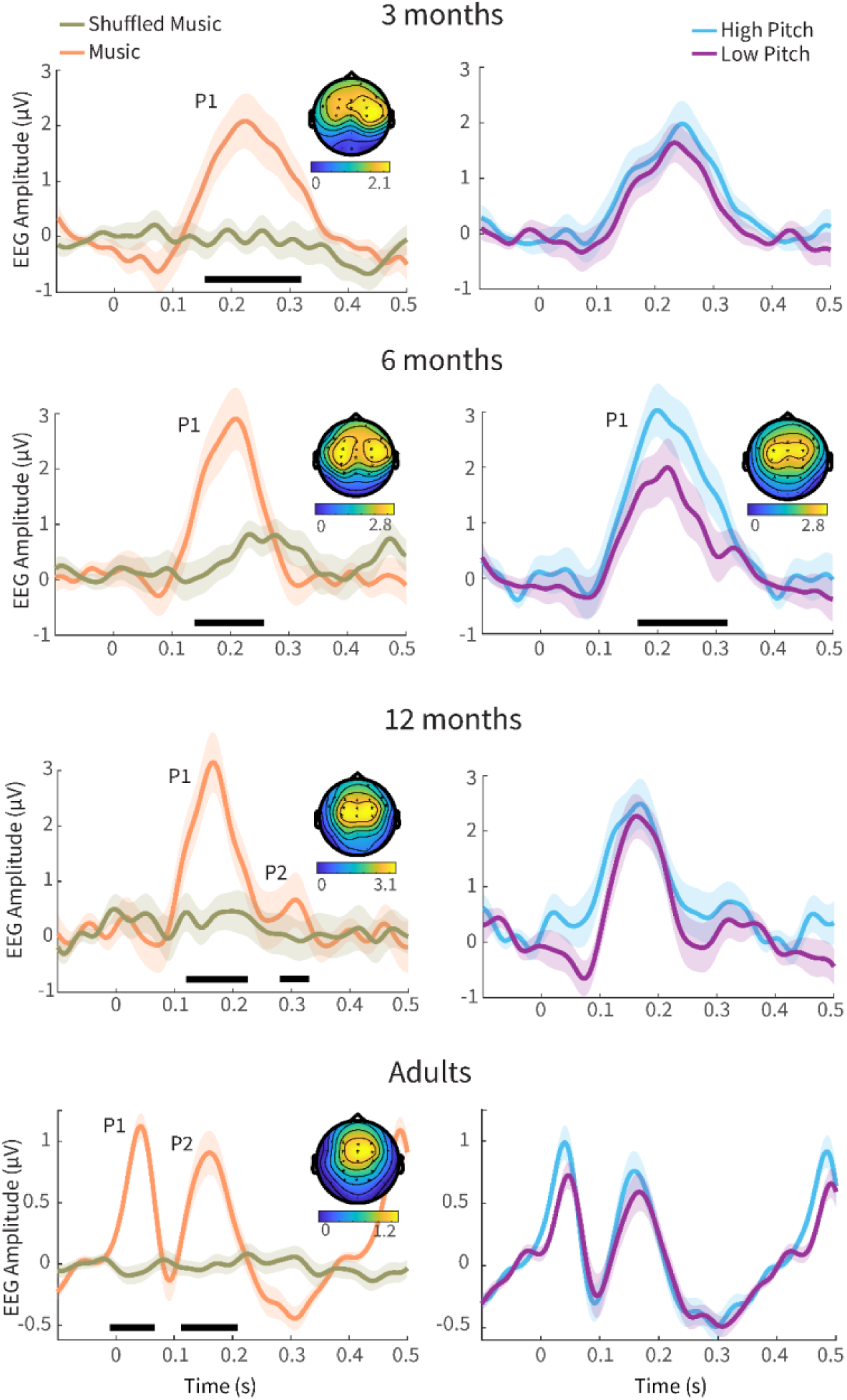
Event-related potentials (ERPs) elicited by the notes comprised within the music (orange, left) vs shuffled music (khaki, left) as well as by the notes comprised within the high-pitch (light blue, right) vs low-pitch music (purple, right), across four groups of participants (plotted in ascending order of age, from top to bottom): 3-, 6-, 12-month-old infants (N=79) and adults (N=26). Grand-average ERPs are averaged across electrodes within the significant cluster of each age group in the music condition (except for pitch condition comparison in the 6-month-olds). Shaded areas indicate the standard error. ERPs show progressively shorter latencies with increasing age. All groups exhibited a P1 response, while only older infants (12-month-olds) and adults additionally exhibited a P2. Music elicited a larger P1 (and, when present, P2) amplitude compared to shuffled music, notably across all groups (time ranges associated with a significant difference are indicated by horizontal black lines). The topography of this neural response (averaged across the time window of the P1 cluster) in the music condition appears to shift more medially with increasing age. Colorbars beneath topography plots index EEG amplitude values.

All infants’ ERPs showed a P1 response, while a P2 response was observed only in 12-month-old infants, albeit with a lower amplitude than the P1. The P1 latency decreased (*Χ^2^*(2)=391.25, *p*<.001), and its amplitude increased (*Χ^2^*(2)=8.59, *p*=.014) with age (*Fig. 2*, left column). Importantly, and in line with the adults’ data, all infant groups exhibited enhanced P1 amplitudes in response to music compared to shuffled music. Actually, across all groups, shuffled music did not elicit clear ERPs as the ones elicited by music. Cluster-based permutation (*n_Perm_*=1000) testing revealed that 3-month-old infants’ P1 amplitude was enhanced between 177 and 305 ms post-stimulus (*cluster-t*=1111.90, *p*=.002). Within this window, the P1 peaked at 212 ms and reached an average amplitude of 1.8 µV. The topography included a frontocentral cluster with a slight right lateralization. In 6-month-old infants, the amplitude of the P1 was enhanced between 116 and 284 ms post-stimulus (*cluster-t*=1401.60, *p*=.002), peaking at 165 ms and reaching an amplitude of 2.8 µV. The topography included a few (centro-) parietal electrodes in addition to several frontocentral electrodes, with a bilateral activation. In 12-month-old infants, the amplitude of the neural response to music was enhanced in a two-peak cluster (*cluster-t*=1416.30, *p*=.002). The first peak, an infantile P1, occurred between 104 and 227 ms, peaked at 146 ms post-stimulus and reached an amplitude of 3.1 µV. Notably, 12-month-old infants exhibited an additional positivity, namely an infantile P2, possibly homologous to the P200 observed in adults. The P2 ranged from 307 to 325 ms post-stimulus and peaked at 316 ms, with an average amplitude of 1.026 µV. The topographies remained frontocentral but were more medial, similar to adults.

To rule out the possibility that increased neural responses in the music condition reflected differences in inter-tone temporal spacing (see IOI range in *Table 1*), we performed control analyses including only epochs from the shuffled music condition for which the prior or subsequent IOI exceeded the median IOI duration. The resulting ERPs in the shuffled music condition were highly similar to those reported above (compare *Fig. 2* with *Fig. S1*), and the difference between the music and shuffled music conditions remained significant across all age groups (*p*<.05). These findings indicate that the observed effects are not solely attributable to variations in inter-tone spacing but also reflect sensitivity to musical structure.

Next, we examined neural responses to the notes in the high- and low-pitch conditions (*Fig. 2,* right column). The morphology of both adults’ and infants’ ERPs was generally comparable to that elicited by the music condition. Cluster-based permutation (*n_Perm_*=1000) testing revealed that the amplitude of the adults’ ERPs was comparable across high- and low-pitch conditions (*p*>.050). This was also the case for both 3- and 12-month-old infants, but notably not for 6-month-old infants, who exhibited an enhanced P1 in response to high-pitch vs low-pitch conditions (*cluster-t*=763.84, *p*=.002). This enhanced positivity (178 and 332 ms) peaked at 204 ms and reached an average amplitude of 2.8 µV. Similar to the neural response elicited by the music condition, the topography included few (centro-) parietal electrodes in addition to several frontocentral electrodes, with a bilateral activation.

### EEG: Auditory Steady State Responses (ASSR)

*Figure 3* shows bar plots indexing the relative power of ASSRs elicited by the auditory stimuli (power estimates were averaged across the electrodes comprised within the ERP clusters that were common to all age groups, i.e., FP2, F7, F3, Fz, F4, F8, FC7, FC3, FCz, FC4, FC8, C3, Cz, C4; see *Fig. 2*).

**Figure 3.**
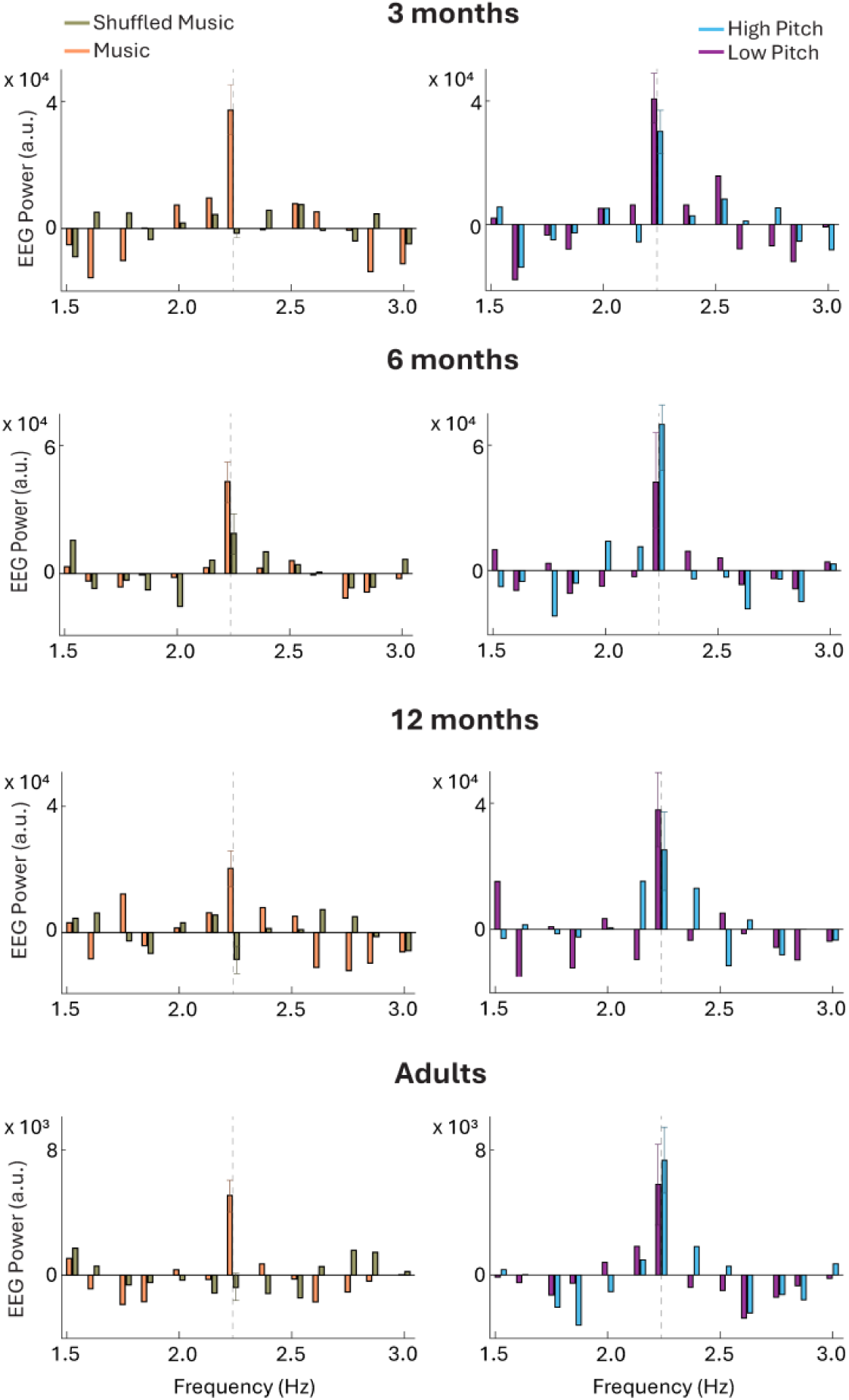
Relative EEG Power (arbitrary unit [a.u.], y-axis) of the auditory steady-state responses (ASSR) elicited by music versus shuffled music (orange and khaki, left), and high-pitch versus low-pitch musical stimuli (blue and purple, right), across four groups of listeners: 3-month-olds (first row), 6-month-olds (second row), 12-month-olds (third row), and adults (fourth row). ASSR power estimates at the frequency (x-axis) matching the musical beat (2.25 Hz, highlighted by vertical dashed lines and including standard error bars) were statistically higher when elicited by music compared to shuffled music across nearly all participant groups (i.e., all but 6-month-olds). High- and low-pitch stimuli evoked similar ASSR (at 2.25 Hz). These results generally align with the ERP results (Fig. 2) across most infant groups and adults, except for 6-month-old infants for whom differences across conditions were either trending (music vs shuffled) or not significant (high vs low pitch).

Our frequency of interest was 2.25 Hz, matching the musical beat as well as the presentation rate of the majority of the notes (15 out of 16 notes) in the auditory sequences (see methods, section “Stimuli”). We used linear mixed models, including power estimates as the dependent variable to contrast the different conditions and age groups (fixed and interaction effects). Participants were modelled as random intercepts. Power estimates were generally higher in response to music as opposed to shuffled music. Model outputs indicated that the power of the ASSR elicited by the music condition was significantly higher than that elicited by the shuffled music condition in 3-month-old infants (*F*(1,50)=7.82, *p*=.007), 12-month-old infants (*F*(1,52)=12.03, *p*=.001) and adults (*F*(1,50)=13.49, *p*<.001); while it only reached a trend for significance in 6-month-old infants (*F*(1,50)=2.95, *p*=.092). ASSR power estimates were not different across high-pitch and low-pitch conditions across all age groups (*p*>.240). These results indicated that most groups tracked the periodicity of the musical beat, leading to an enhancement of power in ASSR at a frequency matching the musical beat. This is generally in line with the ERP results showing stronger neural responses to music than shuffled music (even though 6-month-old infants only showed a marginal difference).

### Extraction of Principal Movements and Estimation of Quantity of Movement

Using principal component analysis, we decomposed full-body kinematics into 10 Principal Movements (PMs), explaining 79.7 % of the whole kinematic variance. The PMs (depicted in Fig. 4) were reminiscent of common infant movements such as front-back rocking (PM1), side sway (PM2), proto-clapping (PM3), leg kicking (PM4), up-down rocking (PM5), arm pedalling (PM6), feet kicking (PM7), whole body wiggling (PM8), feet shuffling (PM9) and feet pedalling (PM10). Labels were assigned qualitatively, following visual inspection of the PMs.

**Figure 4.**
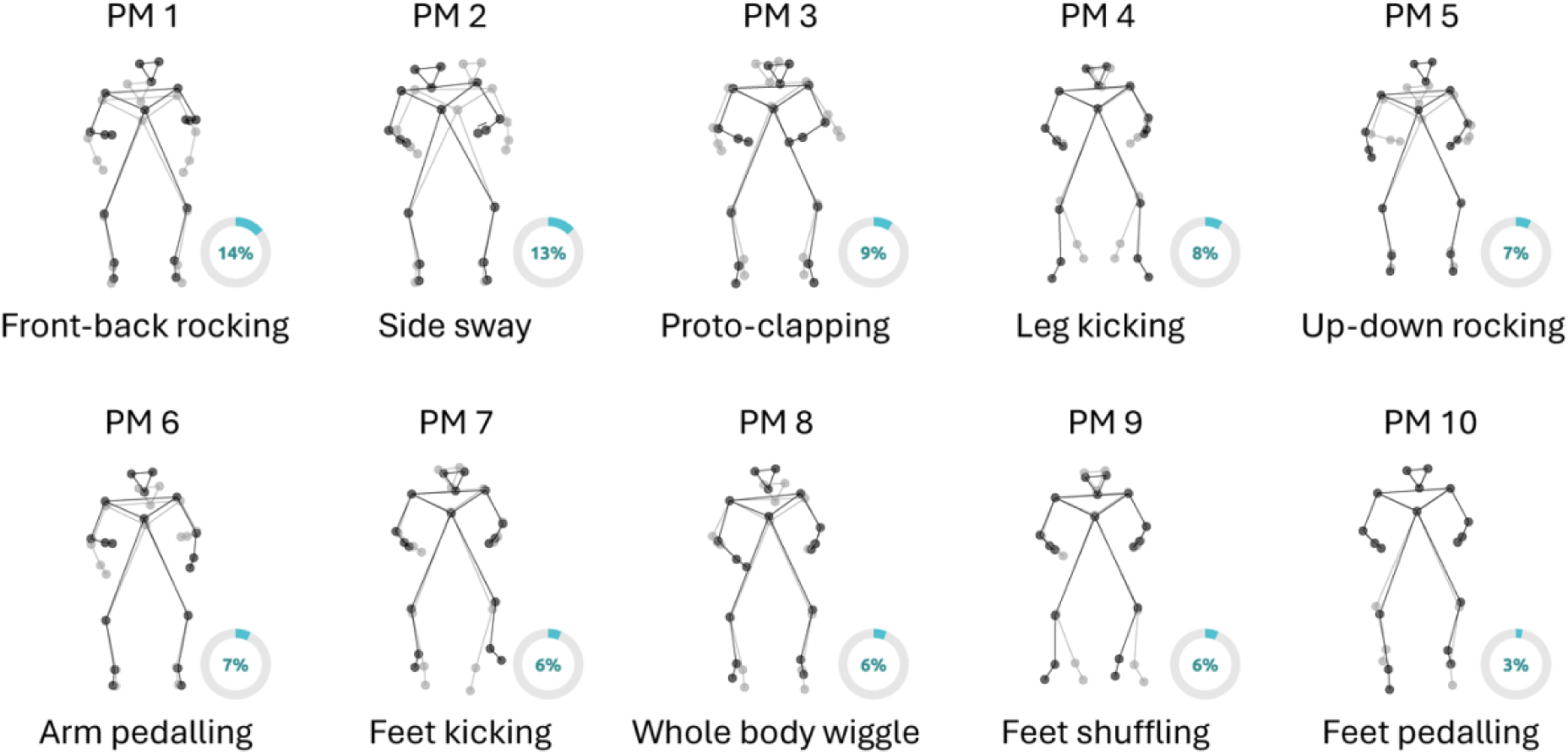
Infants’ principal movements (PMs). PMs are illustrated by showing the two most different body postures (min and max of the PM score, in grey and black, respectively) from the frontal perspective. The reader should interpret the PM as the kinematic displacement necessary to shift from one body posture (grey) to the other (black). Circle diagrams denote the proportion (%) of kinematic variance explained by each PM. Together, the ten PMs account for 79.7% of the total kinematic variance.

Quantity of Movement (QoM) was estimated and compared across PMs, conditions, and groups (*Fig. 5*). QoM estimates were based on first-degree differentiation of the PM time series (see methods). A random intercept was included for each infant. Linear mixed effect modelling yielded a significant interaction between Condition and Age (χ²(2)=16.76, *p*<.001) indicating that only 12-month-old infants exhibited higher QoM in response to music compared to shuffled music across all PMs (i.e., when QoM was computed as an average across all PMs). Post-hoc contrasts comparing QoM averages between conditions revealed a highly significant effect (*t(69.8)*=4.86, *p*<.001; see top left panel of Fig. 5 for QoM averaged across PMs).

**Figure 5.**
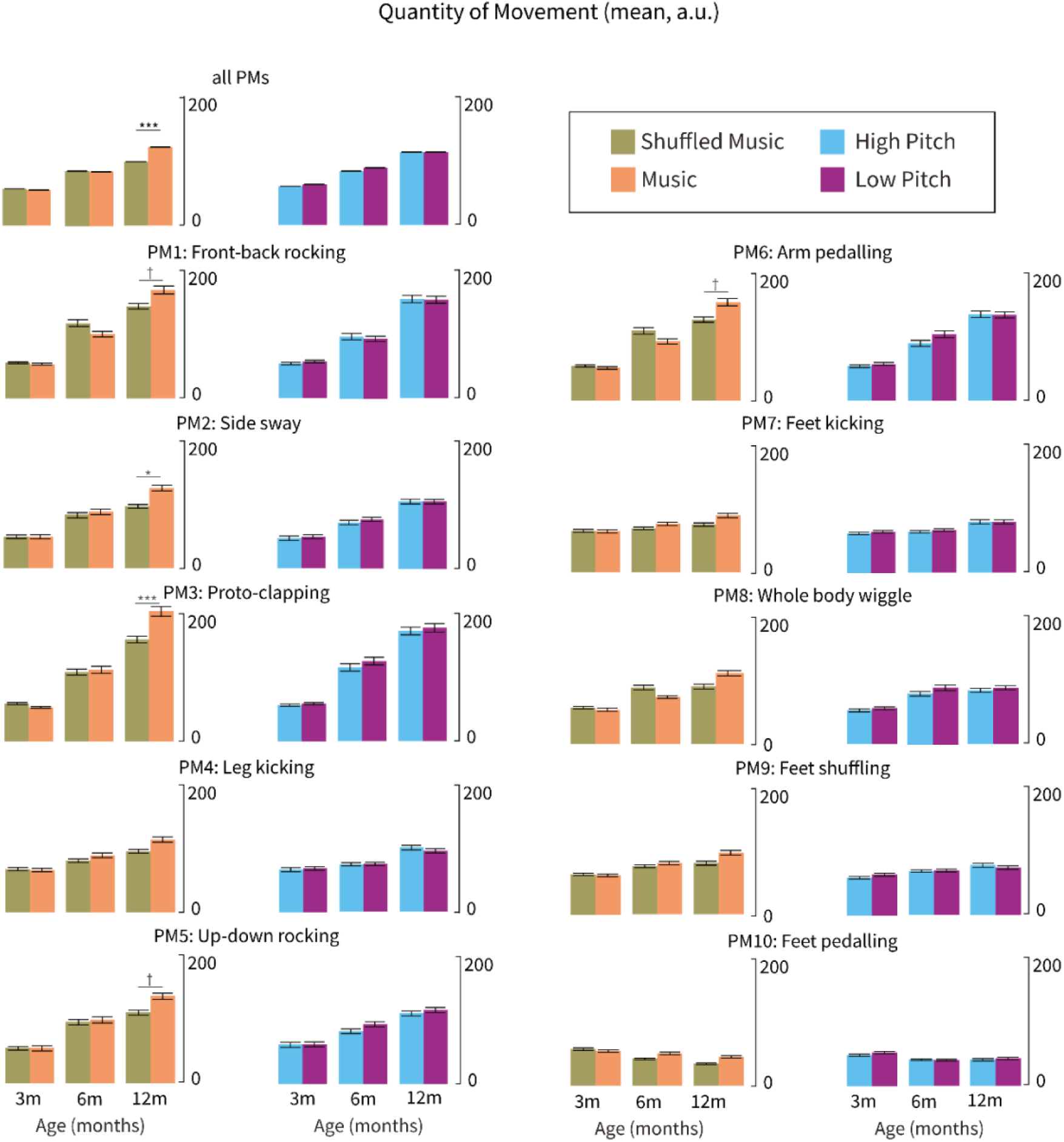
Quantity of movement (mean, a.u.; [QoM]) elicited by music (orange) versus shuffled music (khaki) and high-pitch (blue) versus low-pitch music (purple) across different age groups (3-month-olds, 6-month-olds, 12-month-olds) and principal movements (PMs). Bar plots indicate the mean and standard error of QoM across different age groups, conditions, and PMs. Only twelve-month-old infants showed significantly increased QoM in response to music compared to shuffled music, specifically in PMs involving upper body movements (front-back rocking, side sway, proto-clapping, up-down rocking, and arm pedalling). No significant differences were observed between high- and low-pitch conditions. These results were also replicated in a supplementary analysis assessing differences in variance of (as opposed to mean) QoM (see Fig. S2 and Supplements for more details). † = p<.100, * = p<.050, ** = p<.010, *** = p<.001

Even though there was no interaction effect between Condition, Age and PMs, we still ran post-hoc comparisons to gain preliminary evidence about specific PMs driving the above-described effects. Results indicated that differences in 12-month-olds’ QoM in response to music vs shuffled music were mostly driven by movements of the upper body and/or upper limbs. Specifically, front-back rocking (PM1), side sway (PM2), proto-clapping (PM3), up-down rocking (PM5) and arm pedalling (PM6) were linked with significantly higher QoM in response to music as opposed to shuffled music (PMs 1,2,3,5,6; *ps*<.050, corrected using the false-discovery rate). Contrarily, younger infants (3- and 6-month-olds) did not exhibit significantly different QoM in response to music vs shuffled music in any of the PMs (*ps*>.123). Further, when comparing infants’ QoM to music at different pitches, we found no significant differences between conditions across PMs and age groups (*ps*>.295).

The model also yielded a significant interaction between PMs and Age (χ²(18)=181.575, *p*<.001), indicating that QoM generally increased with age but differently across PMs. Specifically, from 3 to 6 months, the PMs associated with a faster increase in QoM were front-back rocking (PM1), proto-clapping (PM3) and arm pedalling (PM6) (*t*(95.4)=2.34-2.60, *p*=.016-.032). From 3 to 12 months, front-back rocking (PM1), side sway (PM2), proto-clapping (PM3), up-down rocking (PM5) and arm pedalling (PM6) became more prevalent (*t*(95.4)=2.69-5.06, *p*=.001-.025). From 6 to 12 months, proto-clapping (PM3) became even more prevalent (*t*(95.4)=2.71, *p*=.012). Across the first year of life, infants seemed to consistently move their lower body while slowly increasing their capacity for upper-body and whole-body movements while seated.

### Movement: Granger Causality Analysis

Beyond looking at how much infants moved, we further investigated whether the spontaneous occurrence of infant movements could be explained by preceding changes in the intensity of the auditory stimuli. To do so, we used the sound envelope of the auditory stimuli (indexing changes in intensity over time) to predict infant movement velocity (time series representing changes in movement velocity, averaged across all PMs) and vice versa, using Granger-Causality analysis. Prediction estimates (Granger F-values) were yielded across different time lags, indexing the elapsed time necessary for a change in stimulus intensity to predict a change in movement velocity (i.e., highest F-values represent optimal prediction).

A preliminary analysis (sanity check) showed that musical stimuli predicted subsequent movement velocity better than vice versa across all age groups (*Fig. 6*, top-right corner; *F*(1)=94.62, *p*<.001). To identify the optimal temporal lags predicting sound-driven changes in movement, we conducted bootstrapped t-tests corrected for multiple comparisons across time points. Results (*Fig. 6*, left) showed that musical stimuli were better predictors of movement than shuffled stimuli at 3 months (lags of 160-200 ms, *t*(40)=3.55-3.98, *p*<.001), 6 months (lags of 120-240 ms, *t*(37)=2.99-4.49, *p*<=.001) and 12 months (lags of 160-240 ms, *t*(31.6)=4.95-5.03, *p*<.001). Conversely, there was a drop in prediction at later time lags in all groups (∼320-360 ms; *t*(43.6)=3.88-8.25, *p*<.001). Together, these results suggest that infants’ movement was related to intensity changes in the music, but not in the shuffled music.

**Figure 6.**
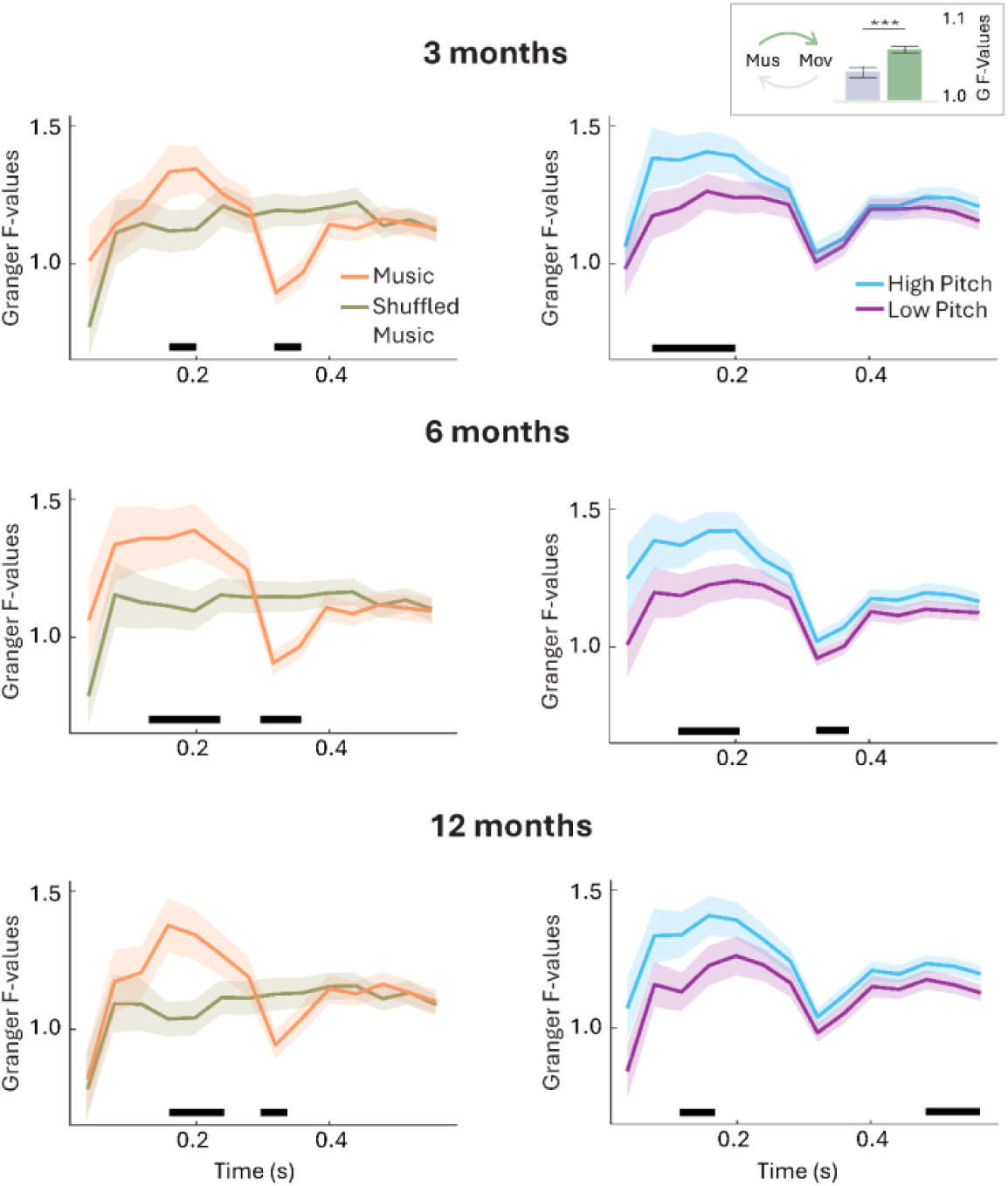
Music-driven movement (Granger-Causality analysis). Top-right: A sanity check analysis showed that musical stimuli predicted subsequent movement velocity (green) better than vice versa (grey; p<.001). Left: Movement velocity was better predicted by music (orange) than by shuffled music (khaki), particularly with time lags of 160-200 ms (shaded areas indicate standard errors; horizontal black lines underline time ranges associated with a significant difference between conditions). Right: Movement was better predicted by high-pitch music (blue) compared to low-pitch music (purple).

Results associated with the high- and low-pitch conditions yielded similar Granger F-values to the music condition (*Fig. 6*, right). Notably, prediction estimates were generally higher for the high-pitch condition as compared to the low-pitch condition, indicating that high-pitch music was a better predictor of movement than low-pitch music. The difference between these two conditions was significant in one time-window in the 3-month-old infants (80-200 ms, *t*(39.26)=2.66-3.81, *p*<.002), while it encompassed two time windows in the 6-month-olds (120-200 ms, *t*(45.64)=3.14-4.21, *p*<=.004, and 320-360 ms, *t*(45.7)=3.32-3.82, *p*<=.004) and 12-month-olds (120-160ms, *t*(40.4)=3.20-3.73, *p*<=.002 and 520-600 ms, *t*(39.9)=3.13-3.41, *p*<.004) – perhaps suggesting that this effect grows stronger with age. Finally, Granger Causality statistics stratified by each PM and age group are detailed in the Supplements and *Fig. S2*.

### Movement: Phase-locked changes and periodicity

The Granger Causality analysis indicated that changes in music intensity drove movement in time, especially at a time lag comprised within a 200 ms delay. This result indirectly suggests that changes in music might evoke a phase-locked movement response. If so, we should be able to observe such phase-locked response when epoching movement data to peaks in the amplitude envelope of the auditory stimuli. To test this prediction, we ran supplementary event-related analyses on the movement velocity time series used for the previous analysis. Cluster-based permutation analyses revealed no significant clusters across age groups and conditions (*ps*>.050; see Supplements, *Fig. S3*), even though it should be noted that 12-month-old infants exhibited a movement peak at ∼200 ms, which was slightly but not significantly higher in response to music vs shuffled conditions. These results indicated that infants did not consistently exhibit phase-locked movement responses to musical events such as peaks in the amplitude envelope of the auditory stimuli.

Next, we also examined to what extent spontaneous movements were periodic and how so across conditions. This analysis builds upon the distribution of the highest coefficients yielded by auto-correlation analyses (similar to Zentner & Eerola, 2010). The results indicated that overall, spontaneous movements tended to be periodic, but such periodicity did not match the musical beat and was not different across conditions (see *Supplements*). In other words, movement periodicity did not result in coordination with music, and it was not modulated by whether infants were listening to music vs shuffled music or high- vs low-pitch music (*Fig. S4*). This result was further supported by an exploratory analysis in which we identified movement bursts by isolating epochs with high amplitude variation in the time series and excluding segments with low or no variation (c.f. Fujii et al., 2014). Together, these results suggest that while infants might generally exhibit rhythmic movements in response to sounds, the specificity of this behavioural response to music – and its coordination with musical structure – continues to develop beyond 12 months of age.

## Discussion

This study examined the development of infants’ neural and movement responses to music during the first year of life using a combination of neural measures, such as ERP and ASSR, alongside quantitative analyses of infants’ movement. These approaches allowed us to explore both sensory and motor components of musicality, shedding light on how infants process and respond to musical stimuli. Below, we discuss 1) the development of neural auditory responses to music, 2) the emergence of music-driven movement patterns, and 3) the sensitivity of both components to pitch across early development.

### Neural Responses to Music: Sensitivity to structured music across all ages

Much research suggests that the infants’ auditory system is sensitive to structural features of music, such as timing and pitch regularities, from early on (Saffran et al., 1999; Thiessen & Saffran, 2009; see Nguyen et al., 2023 for a review). However, the developmental trajectory of such musical sensitivity across the first year of life is still underexplored. Here, we provide a detailed characterization of the progressive maturation of auditory evoked potentials, neurophysiological measures that generally capture rapid changes in the sensory environment, such as the onset of musical notes (Kushnerenko et al., 2002; Somervail et al., 2021). Specifically, we observed ERPs with progressively shorter latency and larger amplitude throughout the first year of life. While these findings likely reflect a general maturation of sensory systems, they also highlight how the evoked potentials elicited by musical stimuli were enhanced in amplitude compared to those elicited by shuffled (i.e., structure-free) stimuli, an effect consistently observed across all age groups. This is in line with the idea that sensitivity to musical regularities emerges early and persists throughout infancy (Trevarthen, 1999). Indeed, sensitivity to simple rhythmic or pitch regularities has been observed in infants at various ages (Cirelli et al., 2016; Edalati et al., 2023; Flaten et al., 2022; Háden et al., 2009, 2024; Stefanics et al., 2009; Winkler et al., 2009), and, in the case of rhythm, even in premature neonates (Saadatmehr et al., 2025). Such sensitivity may be rooted in general auditory predictive processing, whereby the brain extracts regularities from past observations to generate and update predictions about incoming sensory information (Friston, 2010; Köster et al., 2020; Vuust & Witek, 2014). Ecologically valid music, such as the refrains used here, naturally incorporates rhythmic and pitch regularities that could trigger the generation of predictions. This process was markedly dampened or interrupted by shuffled music (Bianco et al., 2024, 2025; Lense et al., 2022), a finding that could be interpreted as evidence of disengagement from such highly unpredictable sequences (Dayan et al., 2000; Esber & Haselgrove, 2011; Kidd et al., 2014).

Which specific musical regularities drove infants’ predictions in our study? Previous EEG research suggests that in newborns, probabilistic auditory predictions are primarily driven by timing rather than pitch regularities when listening to ecologically valid music (Bianco et al., 2025). In contrast, adults rely on both timing and pitch regularities to generate predictions (Di Liberto et al., 2020). Given that our study examined infants older than newborns but not yet as mature as adults, it is likely that timing regularities played a key role for the youngest participants, while pitch regularities may have had a greater influence on the older age groups. However, it is important to note a limitation of such interpretation of our findings: our shuffling procedure permuted both the timing and pitch of the tones simultaneously, which prevents us from disentangling whether the stronger neural responses observed for the music condition reflect sensitivity to rhythmic regularity, melodic structure, or their combination. Hence, to clarify this issue, future research should investigate when, during development, the human brain begins incorporating pitch alongside timing regularities to generate musical predictions.

To explore the underlying neural mechanisms driving the above-described neural process, we analysed ASSR besides ERPs. This was done under the assumption that if the described results were driven by neural entrainment to the beat of the music, then ASSR might better capture differences across conditions, as this measure can capture oscillatory activity. Instead, if the results were driven by evoked responses, then the ASSR results would generally align with ERP results or even be slightly less robust, given that frequency-domain analysis is less suited for capturing time-domain neural modulations. Results showed enhanced ASSR at a frequency matching the musical beat, with stronger power in response to music as opposed to shuffled stimuli and no differences between high- and low-pitch musical stimuli. Hence, ASSR and ERP results were very similar, a conclusion that is in line with previous accounts suggesting that on some occasions, the two measures might originate from the same signal, i.e., evoked responses (Capilla et al., 2011; Novembre & Iannetti, 2018). Further, building on evidence indicating that the infantile P1 originates from the auditory cortex (Chen et al., 2016; Musacchia et al., 2017; Riva et al., 2018), this result sheds light on one of several underlying neural structures that might be responsible for identifying musical structure in the auditory input.

### Motor engagement with Music: Movement Sophistication develops with Age

In adults, music engages not only the auditory system but also the motor system (Fujioka et al., 2012; Grahn & Brett, 2007; Novembre & Keller, 2018; Phillips-Silver & Keller, 2012), often leading to spontaneous motor responses coordinated with the musical input (Hurley et al., 2014; Janata et al., 2012). While some evidence indicates that such spontaneous body movements are also exhibited by infants (Provasi et al., 2014), it remains unclear how this behaviour matures throughout development and how sophisticated it is.

First, as a coarse measure of auditory-motor coupling, we used Granger Causality to predict the time course of body movements using the time course of the auditory input (specifically the amplitude envelope). Taking this approach, we could predict body movements of infants from 3 to 12 months, specifically while listening to music as opposed to the shuffled conditions. This result indicates that recognising musical structures leads not only to distinct neural encoding patterns (as discussed in the previous section) but also to different levels of motor engagement. Further, as this result was observed across all ages, we might speculate that such audio-motor coupling, specifically triggered by music, emerges early in development and might be biologically predisposed. This conclusion aligns with previous findings showing that even foetuses as young as 28–35 weeks gestational age exhibit movement responses to musical sounds (Kisilevsky et al., 2014). However, it is important to note that these prenatal motor responses were not compared to motor responses to other auditory stimuli, as in our study, therefore raising questions about their specificity.

Next, taking a more complex measure of movement, we used principal component analysis to break down body kinematics into 10 independent principal movements (PMs; <u>Bigand et al., 2024; Toiviainen et al., 2010)</u> that explained nearly 80% of the whole kinematic variance. To the best of our knowledge, this approach has never been adopted for the analysis of infants’ datasets, which normally do not distinguish between different kinds of movements (Fujii et al., 2014; Nguyen, Reisner, et al., 2023; Zentner & Eerola, 2010) or do so qualitatively (Thelen 1979, 1981). Building on this data-driven approach, we compared to what extent different movements were exhibited by infants across different ages and conditions. We found that only 12-month-old infants exhibited more movement in response to music compared to shuffled music. This result was driven by specific upper-body movements such as front-back rocking, side swaying, proto-clapping, and arm pedalling. It is not straightforward to interpret why these (and not other) movements were triggered by music, and why only at this age. Potential explanations include the development of refined postural control, typically achieved by 9–10 months (Hadders-Algra, 2005), and the fact that infants were tested in a seated position (a measure taken to simplify comparability across age groups). This seating arrangement likely facilitated upper body movements as the feet were resting. Further studies are needed to systematically characterize infants’ movements to music across different contexts.

Finally, we examined movement coordination – a potential precursor of dance – and tested whether the periodicity of spontaneous movements matched the periodicity of the music. We did not find any evidence of movement coordination in none of the age groups. This result is in line with previous studies (Fujii et al., 2014; Zentner & Eerola, 2010) and suggests that sensorimotor transformation of music, specifically the ability to preserve its periodicity, develops after the first year of life. This delayed emergence may be due to motor control skills required for such transformation, which indeed are known to continue to mature during toddlerhood and even middle childhood (Kim & Schachner, 2022; Phillips-Silver et al., 2024). Hence, while coarse auditory-motor coupling is present at all ages, diversified movement patterns emerge only by 12 months, and spontaneous coordination with music likely continues to develop throughout infancy into childhood. Taking a broader perspective on infants’ motor development, our findings align with research on locomotion across the first 14 months of life, which shows that as the number of motor primitives increases, their intrinsic variability decreases (Hinnekens et al., 2023). Viewed together, these patterns point toward a gradual refinement of motor control: the human motor system first develops the capacity to control individual muscles, and gradually the capacity to integrate them into motor synergies that support complex, coordinated behaviours, such as locomotion, musical synchronization, and dance.

We suggest that the increasing complexity of infants’ motor response to music is linked to the gradual maturation of the dorsal auditory stream, which connects posterior regions of the superior temporal gyrus with premotor cortices (e.g., <u>Chen et al., 2009; Grahn</u> <u>& Brett, 2007; Kotz et al., 2018)</u>. Although this pathway is present at birth (Friederici, 2011; Perani et al., 2010), it has been suggested to play a crucial role in rhythmic entrainment and beat perception, functions that likely develop further as the pathway matures (Honing, 2018; Merchant & Honing, 2014; Patel & Iversen, 2014). This developmental trajectory aligns with neuroimaging evidence showing that while the ventral linguistic pathway (connecting temporal and frontal regions via the extreme capsule) is well-established at birth, the dorsal pathway—particularly the arcuate fasciculus connecting temporal regions to inferior frontal areas—continues maturing throughout the first postnatal months, with different maturational timelines for dorsal versus ventral connections (Dubois et al., 2016). Our suggestion is indirectly supported by comparative work on non-human primates, whose dorsal auditory stream is less developed than in humans (Merchant & Honing, 2014; Patel & Iversen, 2014). Indeed, research suggests that adult macaques struggle to recognise musical beats (Honing et al., 2012, 2018), while adult chimpanzees—though capable of adjusting their rhythmic sway periodicity in response to the beat (Hattori, 2021; Hattori & Tomonaga, 2020)—are unable to match it accurately, much like infants in our study and previous literature (Zentner & Eerola, 2010).

### Pitch Sensitivity: Auditory and Movement preferences for High-Pitched music

At 6 months, infants exhibited enhanced neural responses to high-pitch music compared to low-pitch music, a specificity not observed at 12 months or in adults. This transient enhancement may reflect a critical period of heightened auditory plasticity, potentially supporting the detection of socially relevant stimuli (Fernald & Kuhl, 1987; Kuhl, 2010). Indeed, high-pitched sounds are prominent in infant-directed speech and singing, which are integral to caregiver-infant interactions and social communication during early infancy (Fernald & Kuhl, 1987; Hilton et al., 2022; Nakata & Trehub, 2011; Trainor & Zacharias, 1998; Tsang & Conrad, 2010). At six months, caregiver-infant face-to-face interactions peak (Beebe et al., 2016; Feldman, 2007), with infants relying more on behavioural cues, often conveyed through auditory modalities (Gratier et al., 2015; Nguyen, Zimmer, et al., 2023), and less on physical objects than in later developmental stages. Thus, the enhanced sensitivity to high-pitched music at 6 months may reflect an essential phase in infants’ developing abilities to process both musical and communicative signals (Shenfield et al., 2003). Toward the end of the first year, attentional focus widens (Cooper et al., 1997; Newman & Hussain, 2006) and face-to-face interactions decrease steeply (Jayaraman et al., 2015). At this stage, strengthening of lower-pitch processing likely leads to more balanced encoding of high- and low-pitch music by 12 months and beyond.

It should also be noted that our musical stimuli comprised polyphonic (two-voice) music, carrying sound frequencies falling within the typical range of infant-directed song (∼200-400 Hz, Cirelli et al., 2020; Nguyen, Reisner, et al., 2023; Trainor & Zacharias, 1998). As such, our results might specifically speak for infants’ ability to separate (and prioritize among) simultaneous communicative auditory streams (Marie & Trainor, 2013; Trainor, 2015). Indeed, other studies presenting one-voice pure tone sequences (single isochronous and isotonous tones) with high vs. low pitch - notably at frequencies outside our range (130 vs. 1237 Hz) - have reported stronger neural responses to relatively low frequencies (Lenc et al., 2023). Together, these contrasting observations suggest that pitch prioritization changes not only throughout development but also depends on the polyphonic complexity and spectral characteristics of the perceived stimuli. Further research might investigate this interesting issue further.

Moving to the movement results, it remains difficult to explain why high-pitch music was generally associated with enhanced auditory-motor coupling compared to low-pitch music. One possibility is that high-pitch music led to higher arousal, akin to infant-directed songs (Cirelli et al., 2020; Juslin & Laukka, 2003), which in turn strengthened the coupling between spontaneous movements and music. Alternatively, or complementarily, low-pitch music might have led to lower arousal or generally reduced attentional processes, thereby weakening auditory-motor coupling. If increased arousal were to result in greater overall movement, we would expect higher movement levels in the high pitch condition; however, this was not observed. QoM analyses based on the PMs did not reveal significant differences between the high pitch and low pitch conditions. This discrepancy may arise because Granger causality captures subtler temporal coupling between movement and music rather than gross movement quantity. Thus, high-pitch music may modulate the timing and coordination of motor responses without necessarily increasing the overall amount of movement. In line with prior work (e.g., Bigand et al., 2024), this interpretation emphasizes that musical coordination often involves changes in coupling strength rather than movement quantity per se. Either way, it is difficult to reconcile the movement results with the ERP results showing enhanced auditory processing of high-pitch music only at 6 months. Perhaps, as infants mature, their sensitivity to high-pitched sounds may integrate with not only broader auditory but also motor processing mechanisms. Such a developmental trajectory might explain the reduced neural specificity to high-pitch music by 12 months, alongside a shift toward more coordinated motor responses to lower-pitched stimuli in adulthood (Cameron et al., 2022; Hove et al., 2014; Stupacher et al., 2016). These speculations could be addressed by future research exploring how musical pitch and rhythm interact to drive movement beyond infancy into adulthood to particularly examine *when* neural and motor engagement with low-pitch music becomes more prominent (Cameron et al., 2022; Stupacher et al., 2013, 2016).

## Conclusion

Our study demonstrates that while robust auditory processing of music is present as early as 3 months, the translation of these sensory processes into organised motor behaviours unfolds gradually over and beyond the first year of life. Specifically, while coarse auditory-motor coupling is present at all ages, diversified movement patterns emerge only by 12 months, and spontaneous coordination with music likely develops throughout infancy and into childhood. Hence, our study provides evidence that, much like the auditory encoding of music, the propensity to move in response to music emerges early in development. This may reflect a biological or early-developing predisposition, eventually leading to dance-like behaviour. However, by 12 months, these motor responses remain relatively underdeveloped. Additionally, our study points to a previously unknown association between high-pitch music and auditory-motor coupling, extending the current research focus on infant-directed communication towards motor engagement and spontaneous behaviour. Together, these findings provide initial insights into how the developing brain gradually transforms music into spontaneous movements of increasing complexity. Future research should extend our characterisation of music-induced movement beyond the first year of life and further explore its (to-date mysterious) functional significance (Hoehl et al., 2021; Markova et al., 2019; Nguyen, Flaten, et al., 2023; Wass et al., 2020).

## Materials & Methods

The study was preregistered: https://aspredicted.org/WW3_3TF. Deviations are listed in the Supplement.

### Infant Experiment

Ninety-eight full-term infants, divided into three different age groups (see Fig. 1C), participated in this experiment. Nineteen infants were excluded due to fussiness (n=9) or technical issues (n=10), leading to a 19,6 % attrition rate (in line with infant EEG studies; Hoehl & Wahl, 2012). The remaining 79 infants were either 3 months (n=26, 14 girls, *mean*=113.04 days, *SD*=5.68 days, *range*=98-120 days), 6 months (n=26, 14 girls, *mean*=195.88 days, *SD*=9.46 days, *range*=182-211 days) or 12-13 months of age (n=27, 9 girls, *mean*=380.44 days, *SD*=14.93 days, *range*=361-413 days). All infants were born with a minimum gestational age of 37 weeks, weighed at least 2500 g at birth, and had no known developmental delays, neurological disorders, or hearing impairments. A primary caregiver gave written informed consent and was present during the experiment. The study was approved by the local Ethics committee of the University of Vienna (no. 00645) in line with the Declaration of Helsinki.

### Stimuli

Auditory stimuli were generated by rearranging the refrains of two polyphonic children’s songs (“La Vaca Lola” and “Hopp Juliska”) using Logic Pro X (Apple, Inc.). Stimuli belonged to four distinct conditions (Fig. 1B). The music condition entailed the presentation of the rearranged refrains. The shuffled music condition entailed a disruption of pitch order and timing regularities of the refrains presented in the music condition. Specifically, the original temporal order of the notes was shuffled (disrupting pitch order), and the IOIs were replaced by a new pool of random values uniformly distributed around the original mean IOI ± 30-50% of the distance between the mean and the minimum original IOI (disrupting timing regularities). The bass and melody notes were treated independently in the shuffling process. In addition, the new pool of IOI was not quantized, so the notes did not fall on the beat, thus disrupting the isochrony of the original playsongs. In the high-pitch condition, the melody was shifted one octave higher than in the music condition. In the low-pitch condition, the bassline was shifted one octave lower than in the music condition. Hence, the two voices composing the high-pitch condition were one octave higher than those composing the low-pitch condition. All stimuli were built using the same musical instruments: the melody was played by a flute (“VHS Flute” from Yamaha DX7, Arturia) and the bassline was played by an electric bass (“Liverpool” electric bass from Logic Pro), with almost all bassline notes (15 out of 16) falling on the beat. Importantly, all stimuli had the same duration (21 s), tempo (135 BPM), tonality (C-major) and loudness (which stayed within a range of 1.5 LUFS, a measure that considers the sensitivity of the human auditory system across frequencies).

### Procedure

#### Setup and Task

Infants were seated in an infant highchair equipped with an age-appropriate baby seat facing a computer screen (1 m distance, see Fig. 1A). Parents sat next to the infants (on their right and slightly towards the back). The stimuli were presented free field (∼60 dB SPL) from two audio speakers (Q Acoustics 3020i) placed on either side of the monitor, facing the infants. During the procedure, infants passively listened to a series of auditory stimuli in the following sequence: First, there were 10 seconds of silence, followed by the shuffled music condition. Then, the music, high-pitch and low-pitch conditions were played in random order. After that, there were another 10 seconds of silence, followed by a second round of the music, high-pitch and low-pitch conditions in random order. Finally, the shuffled music condition was played one last time, followed by 10 seconds of silence. The same song was used in all the different conditions within each presentation block. The order of presentation blocks for each song was counterbalanced. Meanwhile, infants were watching a silent movie of blooming flowers, slowly fading in and out (image duration 10 s). Videos were included to keep infants calm and engaged. Only infants who remained calm for at least two presentation blocks were included in the final sample (n=3 were excluded). Stimulus presentation and the synchronization between EEG and video cameras were controlled in Presentation (Version 23.0, Neuro-behavioral System, Berkeley, CA) utilizing triggers sent at the start of the experiment and the beginning of each trial.

##### Electroencephalography (EEG)

We recorded EEG using a Brain Products BrainAmp DC system with 32 active Ag-AgCl electrodes, mounted on infant-sized EEG caps (Acticap) according to the international 10-10 system. The reference electrode was placed at TP9 (left mastoid), and the ground electrode was placed at Fp1. EEG data were sampled at 1000 Hz. Impedances were below 20 kΩ at the beginning of the experiment.

##### Video Recording

Infant body movement was recorded using three video cameras (Axis Communication, Lund) sampling at 25 Hz with a resolution of 1920 x 1080 pixels. The three video recordings, taken from frontal (0°), diagonal (45°) and side view (90°) perspectives, were synchronized in VideoSyncPro (Mangold International). The cameras were positioned at the height of the infants’ heads to capture their whole bodies.

### Adult Control Experiment

Twenty-six healthy young adults (11 female, *mean* = 27.55 years of age, *SD*: 7.09 years, *range*=19-47 years) participated in the experiment. The study was approved by the Regional Ethics Committee of Liguria (no. 794/2021). Participants underwent the same experimental procedure as the infants, including the same auditory stimuli and video material. EEG was recorded with 64 Ag-AgCl electrodes, closely matched to the infant configuration (see Supplements for further details) and sampled at 1024 Hz. Data were down-sampled to 1000 Hz to match the pre-processing and data analysis procedure (detailed below) of the infant study. The experiment was conducted to provide a ground truth for the interpretation of the infants’ EEG data. Therefore, no video data was recorded to track adults’ movements.

### Data Processing

#### EEG pre-processing

Infant EEG data are notoriously noisy as infants may not understand or comply with task instructions. It follows that their behaviour can lead to artefacts in the EEG recordings. To mitigate this issue, we used a combination of open-access denoising algorithms and adopted a fully data-driven pre-processing pipeline that we previously developed to denoise EEG data recorded from awake monkeys (Bianco et al., 2024) and humans dancing (Bigand, Bianco, Abalde, Nguyen, et al., 2024). The pipeline relied on a combination of MATLAB functions developed for toolboxes such as Fieldtrip (Oostenveld et al., 2010) and eeglab (Delorme & Makeig, 2004). Continuous EEG data were band-pass filtered (0.3 Hz - 30 Hz, 3^rd^ order Butterworth filter, zero-phase) and segmented into trials starting 3 seconds before song onset and ending 3 seconds post song offset. Next, noisy and faulty electrodes were identified and rejected by assessing flat-lining for over 5 s (function *clean_flatlines*), correlations between electrodes for *r*<.1 or line noise above 20 SD from the mean of all electrodes (function *clean_channels)*. These initial steps were conducted to find a rough indication of electrodes carrying low-quality signals. In addition, mean, standard deviation, and peak-to-peak distances were calculated over time for each electrode. If any of those variables exceeded 2.5 SD from the mean of the other electrodes, that electrode was (provisionally) discarded. This process was iterated without the outlier electrode(s) until a distribution without outliers was found. Electrodes were then re-referenced to the averaged mastoids (TP9, TP10) or the left mastoid only in cases when TP10 was noisy (N=26). We then used artifact subspace reconstruction (ASR, threshold 5) to remove artefactual EEG activity (Kothe & Jung, 2015). An automatic independent component analysis (ICA) procedure using ICLabel was implemented, and components that were classified as eye components with a minimum probability of 50 % were rejected (mean=0.76, SD=0.67, 0-2 components per infant). Next, previously excluded electrodes were interpolated by using the neighbouring electrodes (spherical spline; mean number of interpolated channels=6, SD=3.09). We assessed data quality (including trial counts and signal-to-noise ratio [SNR]) across age groups and confirmed that these metrics were comparable (see Supplementary Materials for details).

#### Video data pre-processing

We extracted infants’ movement time series by using DeepLabCut (version 2.2.3; Mathis et al., 2018; Nath et al., 2019) in Python on videos filming the infant from the front (see *Fig. 1*). A trained coder (T.N.) labelled 18 body parts per individual (left eye, right eye, mouth, left shoulder, right shoulder, chest, left elbow, left wrist, left hand, right elbow, right wrist, right hand, left knee, left ankle, left foot, right knee, right ankle, right foot) in 10 frames randomly taken from the videos of each infant in each age group (n=24 [3m], 25 [6m] and 24 [12m]; 80% were used for training). We used a ResNet-50 neural network with default parameters for single subject detection for around 5-17 training iterations (correcting labels of 5 frames per video of age groups) until test error plateaued. The test error was 8.27 (3-month-olds), 10.19 (6-month-olds), and 11.89 (12-month-olds) pixels (image size was 1920 by 1080 pixels). The networks of each age group were used to analyse videos from the same age group. The (x and y) coordinates were then pre-processed using a custom code written in Python. Coordinates that were estimated with likelihood values 2 SD away from the mean probability per age group were set to NaN (i.e., not a number). These gaps in the coordinates of body parts were then recovered from predefined neighbouring body parts in a forward and backward approach (from head to feet, followed by feet to head). For example, if the left wrist was missing at t_0_, it was recovered using the position of the left elbow or the left hand at t_0_. Next, we assessed the correctness of the estimated body part displacements, i.e., demeaned coordinates. Specifically, we set the coordinates (either x or y) of each body part below or above 3 SD from the mean of all trials of each infant to NaN (outlier procedure). To further verify left- and right-hand displacements, we checked the distance between the wrist and hand, specifically using the same outlier procedure. The remaining NaN values were replaced using linear interpolation for each body part coordinate in time.

##### Principal movements

We used Principal Component Analysis (PCA) to reduce dimensionality of the pre-processed movement data and at the same time to extract a set of interpretable Principal Movements (PM) that generalize across trials, conditions, and all infants (Bigand, Bianco, Abalde, & Novembre, 2024). To ensure that PCA captured only common movements across participants, rather than postural or anthropometric differences between them, we demeaned and standardized the body part vectors for each trial. Specifically, we subtracted the mean body part vector (averaged over time) from the body part vectors at each frame. Then, these demeaned body part vectors were divided by a global measure of standard deviation across time, which was computed by combining all body parts to preserve the variance differences between body parts. This allowed us to concatenate all trials from all participants into a single data matrix, with each trial contributing equally to the variance of the pooled matrix. PMs were derived from the eigenvectors, and their corresponding scores obtained from the PCA applied to the data matrix:

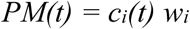

where c_i_(t) is the i^th^ PM score at time t and w**_i_** is the i^th^ PM weight vector, or eigenvector (1 × 36). Each of the PMs reflects the trajectory of covarying body parts. We extracted 10 PMs that explained 79.7 % of the whole kinematic variance (Fig. 3). It is important to note that the PCA does not directly extract “real movements” but rather movement dimensions with large variances. PMs’ time series (score of each principal component) were low-pass filtered at 10 Hz. Next, we calculated the time series’ first-degree derivative to extract velocities and the absolute values of velocities to calculate movement quantities (e.g., Bigand et al., 2024; Górecki & Łuczak, 2013). Velocities include information on movement direction (below or above 0) and are, therefore, particularly useful to analyse correlations with other streams, such as music (detailed in the section on Granger Causality). Movement quantities only consider the amount of movement and can contribute to understanding descriptive differences between conditions, regardless of the movement direction.

### Data analyses

#### EEG: Event-related potentials (ERP) analysis

EEG data were first lowpass filtered at 20 Hz (FIR-type filter) to specifically retain the part of the EEG signal carrying event-related potentials. EEG data were epoched according to the onsets of the bass notes of the musical stimuli (which fell on the beat of the music in 15 out of 16 notes). Epochs included 600 ms of data (−100 ms to +500 ms relative to tone onset – note that notes occurred every 444 ms, so the last 56 ms of each epoch refers to the next tone). In preparation for further analysis, epochs were automatically examined to check if they were still contaminated by artifacts. Epochs were automatically excluded if the standard deviation of the analysed channels exceeded a voltage threshold of 100 μV within a 200 ms interval. All infants and adults contributed at least 165 clean epochs per condition (3m: *M*=340.09, *SD*=138.96; 6m: *M*=373.69, *SD*=144.07; 12m: *M*=361.47, *SD*=157.46; adults: *M*=399.95, *SD*=8.50). Epochs were baseline-corrected by subtracting the average voltage between −100 and 0 ms relative to the tone onset. Epochs belonging to the same experimental condition were averaged together. Event-related EEG amplitude modulations were compared using cluster-based permutation analyses (*n_Perm_*=1000; Maris & Ostenveld, 2007). Clusters were based on both temporal consecutiveness and spatial adjacency of EEG electrodes (3 cm distance). A cluster had to be composed of at least two consecutive time points with a *p-value* <.05 on at least two neighbouring EEG electrodes. The clusters were tested using a *p-value* of <.05 and one-tailed according to the above-detailed hypotheses.

In additional analyses, we extracted the amplitude and latencies of the individual peak of the P1 component. Using two linear mixed-effects models, we tested whether amplitudes (dependent variable) and latencies (dependent variable) were dependent on age and condition (fixed and interaction effect) while assuming a random intercept for each participant.

#### EEG: Auditory Steady State Response (ASSR) analysis

EEG data were segmented according to each condition (21 seconds long, 3 repetitions of the refrain) and averaged over segments of the same condition to enhance the ASSR (see Cirelli et al., 2016). Each segment was submitted to single-taper frequency transformation using a Hilbert taper to estimate frequencies between 0.5 and 5 Hz, with a frequency resolution of 0.25 Hz. Activity unrelated to the frequency of interest was removed by subtracting the average amplitude measured at neighbouring frequency bins (i.e., −5 to −3 bins and +3 to +5 bins). Performing this subtraction removes residual background noise around the frequency bin of interest, leaving only the activity directly related to the ASSR. Next, we extracted amplitudes at the beat-related (2.25 Hz) frequencies and compared these across conditions.

#### Movement: Granger Causality analysis

We conducted a Granger Causality analysis in R (function = *causality,* library = *vars)* to measure to what extent the envelope of the auditory stimuli predicted movement velocity over time and across conditions (see also Klein et al., 2022). To do so, we extracted the broadband amplitude envelope of the auditory stimuli using Hilbert transform. We then conducted Granger Causality analyses on amplitude envelopes predicting the PM velocity time series across different model orders (lags) ranging from 40 ms to 1000 ms. Granger Causality is based on the concept of prediction in the sense that if a time series X_t_ "Granger-causes" another time series Y_t_, then past values of X_t_ contain information that helps predict Y_t_ beyond the information contained in past values of Y_t_ alone. Granger Causality is particularly useful for capturing the temporal relationship between different time series that are characterized by sudden and transient bursts in activity (such as infant rhythmic movement; Thelen, 1979). As a sanity check, we also conducted a reverse analysis predicting music (i.e., the amplitude envelope of the auditory stimuli) from the PM velocity time series and verified that the resulting F-Values (averaged across all lags: 40 – 1000 ms) were statistically lower compared to those yielded by the former (music-to-movement) analysis. Indeed, the sanity check analysis reveals that predictions from audio stimuli to PM velocity time series (*M*=1.08, *SE*=0.009) were significantly higher than from PMs to audio stimuli (*M*=1.03, *SE*=0.009; χ²(1)=1669.2, *p*<.001, see *Fig. 5*).

#### Movement: Event-related analysis

To examine whether the events (i.e., musical notes) comprised within the auditory stimuli would trigger event-related movements, we used an event-related approach (mirroring the ERP analysis discussed above). First, we segmented epochs of 700 ms (between −100 ms and +600 ms relative to peaks in the amplitude). Epochs were then averaged for each condition. Detailed results for this analysis are reported in the Supplements (see *Fig. S2*).

#### Movement: Periodicity analysis

We analysed the movement periodicity using autocorrelation on the same epochs employed in the event-related movement analysis. This means we estimated the autocorrelation of the movement time series in a running window of 3.5 s (length of 1 bar) with 25% overlap, mirroring the approach by Zenter & Eerola (2010). This approach provides an estimation of the periodicity of the PMs. We extracted the local maxima of the autocorrelation values in each window and fitted a normal distribution (Kernel density estimation) to the collection of these values for each condition. We used a bin width of 0.08 and estimated values between 300 and 600 ms according to the beat-related lag in our auditory stimuli. We then compared the beat-related lag (444 ms lag) across conditions. The results are reported in the Supplements (see *Fig. S3*).

### Statistical analyses

Analyses were conducted in R and included non-parametric tests or linear mixed-effects models (LMM; lme4 package). Each participant was modelled as a random intercept in all LMMs. Statistical significance was evaluated by likelihood-ratio tests conducted using *Anova* (car package), and post-hoc contrasts were run using *emmeans* (emmeans package). We controlled for increased Type I error from multiple comparisons, when necessary, using the false-discovery rate (Benjamini & Hochberg, 1995). The statistical significance level was set to α = 0.05.

## Acknowledgements

We are grateful to Eluisa Nimpf, Flavia Arnese, Larissa Reitinger, Josefine Schürholz and Liesbeth Forsthuber for their support in data acquisition, processing, and video coding. Additionally, we thank all families who participated in the study and the Department of Obstetrics and Gynaecology of the Vienna General Hospital for supporting our participant recruitment.

## Credits

**Trinh Nguyen**: Conceptualization, Methodology, Software, Formal Analysis, Investigation, Data curation, Visualization, Writing - Original draft, Writing – Review & Editing, Project administration; **Félix Bigand**: Software, Validation, Data curation, Formal Analysis, Writing – Review & Editing. **Susanne Reisner**: Investigation, Data curation, Writing – Review & Editing, Project administration. **Atesh Koul**: Software, Writing – Review & Editing; **Roberta Bianco**: Validation, Software, Writing - Original draft, Writing – Review & Editing; **Gabriela Markova**: Conceptualization, Data curation, Writing - Review & Editing; **Stefanie Hoehl**: Conceptualization, Methodology, Resources, Writing - Review & Editing, Supervision, Funding acquisition; **Giacomo Novembre**: Conceptualization, Methodology, Resources, Writing - Original draft, Writing - Review & Editing, Supervision, Funding acquisition.

## Data availability

The data and stimuli reported in this manuscript are available in the following repository: https://doi.org/10.48557/DCSCFO. The codes used for analyses and figures are available on Github: https://github.com/tnguyen1992/tinydancer.

