## Supplementary Materials for "Development of Auditory and Spontaneous Movement Responses to Music over the First Postnatal Year"

**Deviations from Pre-Registration**

We ran the majority of analyses according to the preregistration. There is, however, one deviation. Namely, we did not assess motor components (such as mu-oscillations) of infants’ neural responses to music in the current paper. We deemed these analyses too extensive for the current paper and will follow up on these analyses in future studies.

**EEG Electrode Configuration in infants and adults**

Infant electrode configuration included the following electrodes:

VEOG, Fp1 (Ground), Fp2, F7, F3, Fz, F4, F8, FC3, FCz, FC4, FT7, FT8, C3, Cz, C4, T7, T8, CP3, CP4, P7, P3, Pz, P4, P8, TP9 (online Ref), TP10 (offline Ref), PO9, PO10, O1, Oz, O2

The adult electrode configuration included the same electrodes as the infant configuration:

VEOG, Fp1, Fp2, F7, F3, Fz, F4, F8, F4, F8, FC3, FCz, FC4, FT7, FT8, C3, Cz, C4, T7, T8, CP3, CP4, P7, P3, Pz, P4, P8, TP9 (offline Ref), TP10 (offline Ref), O1, Oz, O2. CMS, DRL were used for real-time referencing and grounding.

The following electrodes from the infant configuration were not recorded in the adult configuration: PO9, PO10

The following electrodes were recorded in the adult configuration but excluded in the analysis to make infant and adult electrode configurations comparable: Fpz, AF7, AF3, AFz, AF4, AF8, F5, F1, F2, F6, FC5, FC1, FC2, FC6, C5, C1, C2, C6, TP7, CP5, CP1, CPz, CP2, CP6, P9, P5, P1, P2, P6, P10, PO7, PO3, POz, PO4, PO8, Iz.

**EEG signal quality checks**

We conducted two analyses to compare the EEG data quality across age groups. First, we compared the number of trials that were included in the final analysis per age group. The trial number did not differ significantly across age groups (p > .361). Second, we calculated the SNR by dividing the EEG power at the frequency of interest (i.e., 2.25 Hz, matching the musical beat) by the background noise in surrounding bins (3rd to 5th bin, see ASSR methodology for further details; c.f., Christodoulou et al., 2018; Cirelli et al., 2014). This division yields a signal-to-noise ratio that can be averaged across conditions and compared across age groups to assess variations in signal quality. Here, we find that all three age groups show considerable SNR above 1. Further, even though there were descriptive differences, namely 3- and 6-month-old infants exhibited numerically higher SNR (3m: M=2.569, SD=1.104; 6m: M=2.743, SD=1.001) compared to 12-month-old infants (M=1.907, SD=0.749), the three age groups did not differ significantly (three t-tests, controlled for multiple comparison using the false discovery rate, p > .134). Together, these two analyses indicate that signal quality was comparable across age groups.

*Table S1*. Continued acoustic characteristics of the musical stimuli

| Song name (condition) | *Envelope M±SD (a.u.)* | *Envelope Range (A.u.)* |
| --- | --- | --- |
| Hopp (Music) | 0.162±0.047 | -0.014-0.290 |
| Hopp (Shuffled Music) | 0.161±0.054 | -0.012-0.318 |
| Hopp  (High Pitch) | 0.160±0.046 | -0.011-0.290 |
| Hopp  (Low Pitch) | 0.180±0.046 | -0.007-0.317 |
| Lola  (Music) | 0.147±0.067 | -0.013-0.265 |
| Lola (Shuffled Music) | 0.154±0.059 | -0.005-0.305 |
| Lola  (High Pitch) | 0.144±0.067 | -0.013-0.265 |
| Lola  (Low Pitch) | 0.160±0.069 | -0.011-0.262 |

*Note*. *Hopp indicates the Hungarian playsong (“Hopp Juliska”), Lola indicates the Spanish playsong (“La vaca lola”).*

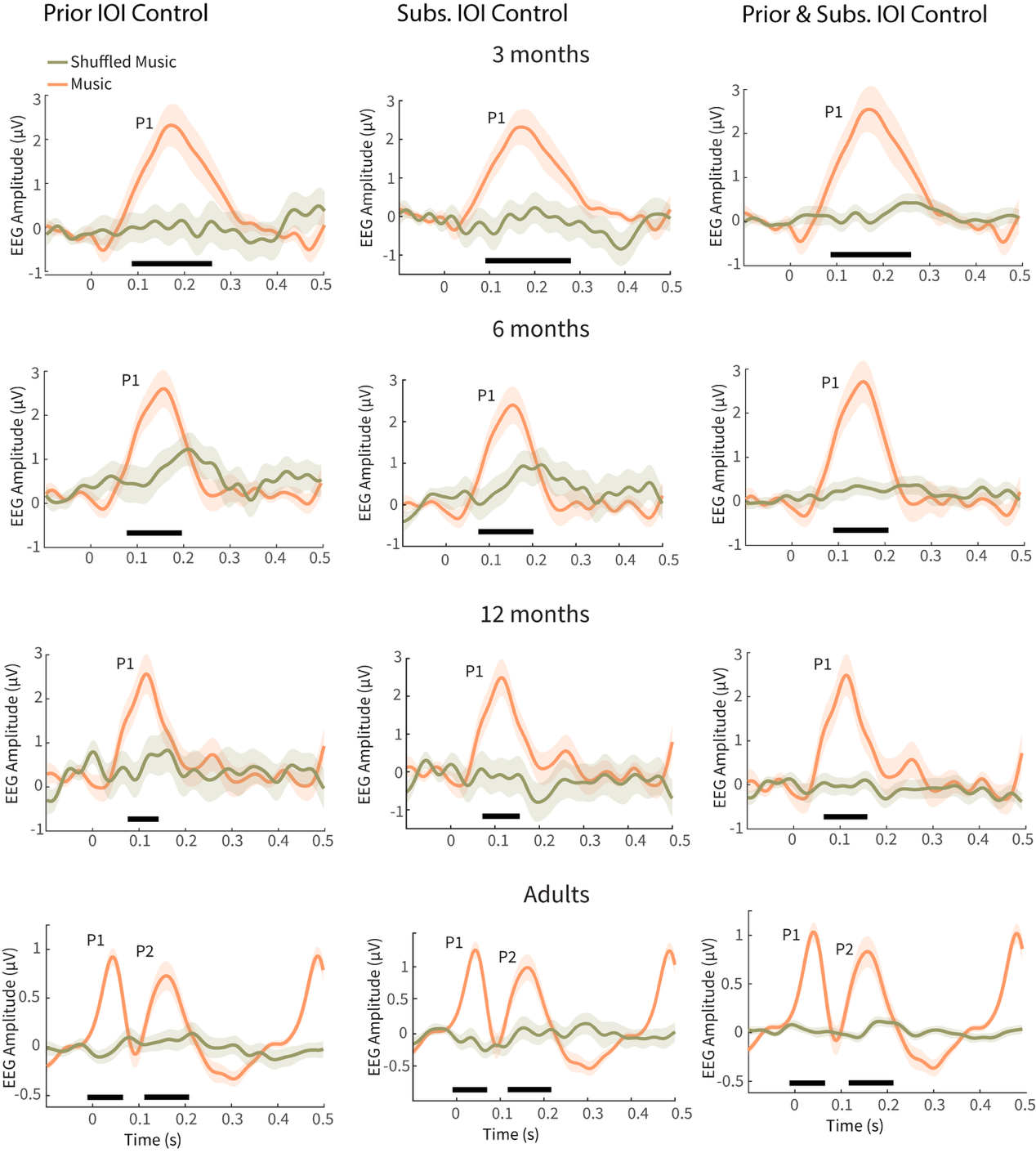

***Figure S1.*** *Event-related potentials (ERPs) elicited by the notes comprised within the music (orange) vs shuffled music (khaki) across four groups of participants from top to bottom: 3-, 6-, 12-month-old infants (N=79) and adults (N=26). These control analyses included only epochs from the shuffled music condition in which the prior (left column), subsequent IOI (middle column) of bassline notes or prior and subsequent IOI of melody and bassline notes (right column) exceeded the median IOI duration (*212 ms*). The number of included epochs in the music condition was matched to the shuffled music condition. Grand-average ERPs were computed by averaging across electrodes within the significant cluster identified for each age group in the music condition based on the primary ERP analysis. Shaded areas indicate the standard error of the mean.*

**Variance in Quantity of Movement**

To assess how infants’ movement patterns varied with musical structure, we analyzed the Variance in Quantity of Movement (QoM) using a mixed-effects model. Model outputs show a significant interaction between age groups and conditions (χ²(2)=8.36, *p*=.015), age groups and PMs ((χ²(18)=242.50, *p*<.001), and main effects of age (χ²(2)=8.66, *p*=.013) and PMs (χ²(9)=288.59, *p*<.001). Post-hoc contrasts indicated that 12‑month‑old infants showed greater variability in QoM when listening to music compared to shuffled music (*t(69.8)*=3.09, *p*=.003). In contrast, 3- and 6-month-olds’ QoM did not vary differently across music and shuffled music (*p*>.591). We also found no significant differences in QoM variance between musical stimuli that differed in pitch (*p*>.339).

**Granger Causality Statistics stratified by each PM and age group**

To assess whether the Granger Causality results were driven by specific PMs, we averaged the Granger F-values within the 160-200 ms lag window and entered these average values into a linear mixed effect model. This analysis yielded an interaction effect between age, PM, and condition (χ²(18)=31.13, *p*=.028, *Fig. S1*), while all other effects were not significant (*p*>.205). Post-hoc contrasts showed that in 3-month-old infants, music vs shuffled music drove mostly particular movements such as up-down rocking (PM5), arm pedaling (PM6), whole body wiggling (PM8), feet shuffling (PM9), and feet pedaling (PM10) (*t*(583)=2.02-3.42, *p*=.001-.044). For 6-month-olds, music drove mostly front-back rocking (PM1), side swaying (PM2), proto-clapping (PM3), feet kicking (PM4), arm pedaling (PM6), and whole body wiggling (PM8) (*t*(573)=2.06-4.13, *p*=.001-.047). In 12-month-olds, music-driven movement mostly entailed front-back rocking (PM1), proto-clapping (PM3), arm pedaling, up-down rocking, feet kicking (PM7), whole body wiggling (PM8), and feet shuffling (PM9) (*t*(573)=1.97-4.87, *p*=.001-.049). Conversely, the same analysis, conducted on high- and low-pitch music conditions, yielded a main effect of condition (χ²(1)=47.87, *p*<.001) indicating that high-pitch music drove movement better than low-pitch music, and that was so across all PMs and age groups (*t*(67.6)=6.87, *p*<.001). Together, these results indicate that the principal movements specifically predicted by music, as opposed to shuffled music, grow in number and change in type with age: in younger infants (3 months), music best predicted lower-body movements, while in 6-month-olds, music best predicted upper-body movements. By 12 months, music best predicted whole-body movements.

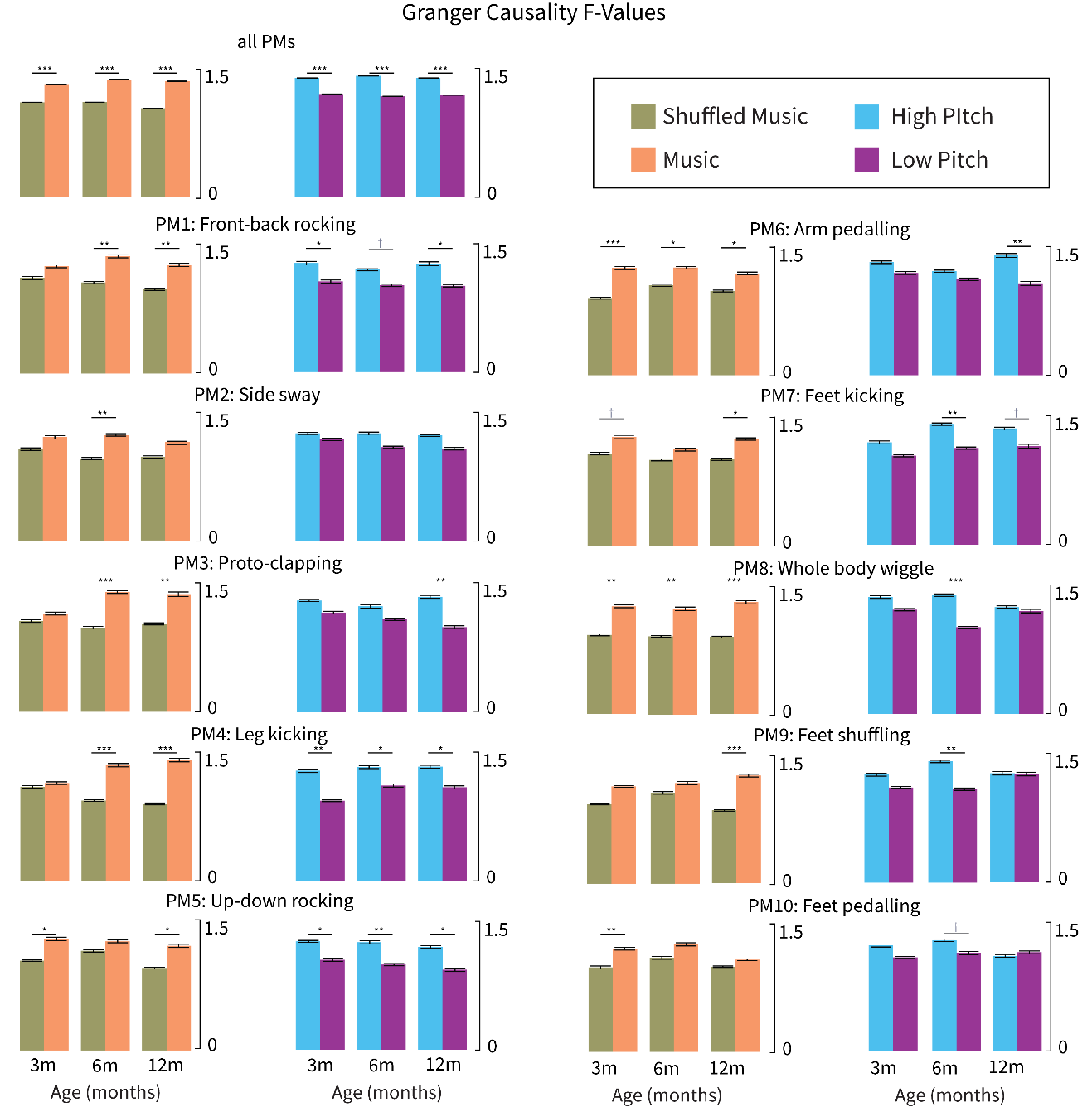

***Figure S2****. The figure displays the Granger Causality F-Values (y Axis) averaged across conditions and age groups for all PMs combined (top row left) as well as for individual PMs (1 to 10, left to right). The left columns include music (orange) vs shuffled music (khaki) contrasts for each age group (x-Axis: 3m, 6m, and 12m) while the right columns include high-pitch (blue) vs low-pitch (purple) contrasts. Across all three age groups movement velocity was more strongly predicted by music than shuffled music, and by high-pitch than low-pitch music, p<.001. These effects were particularly pronounced for movement patterns such as front-back rocking (PM1), proto-clapping (PM3), leg kicking (PM4) and whole-body wiggle (PM8), especially so at 6 and 12 months of age, as reflected in the variation of Granger F-values across PMs and ages.*

**Movement responses**

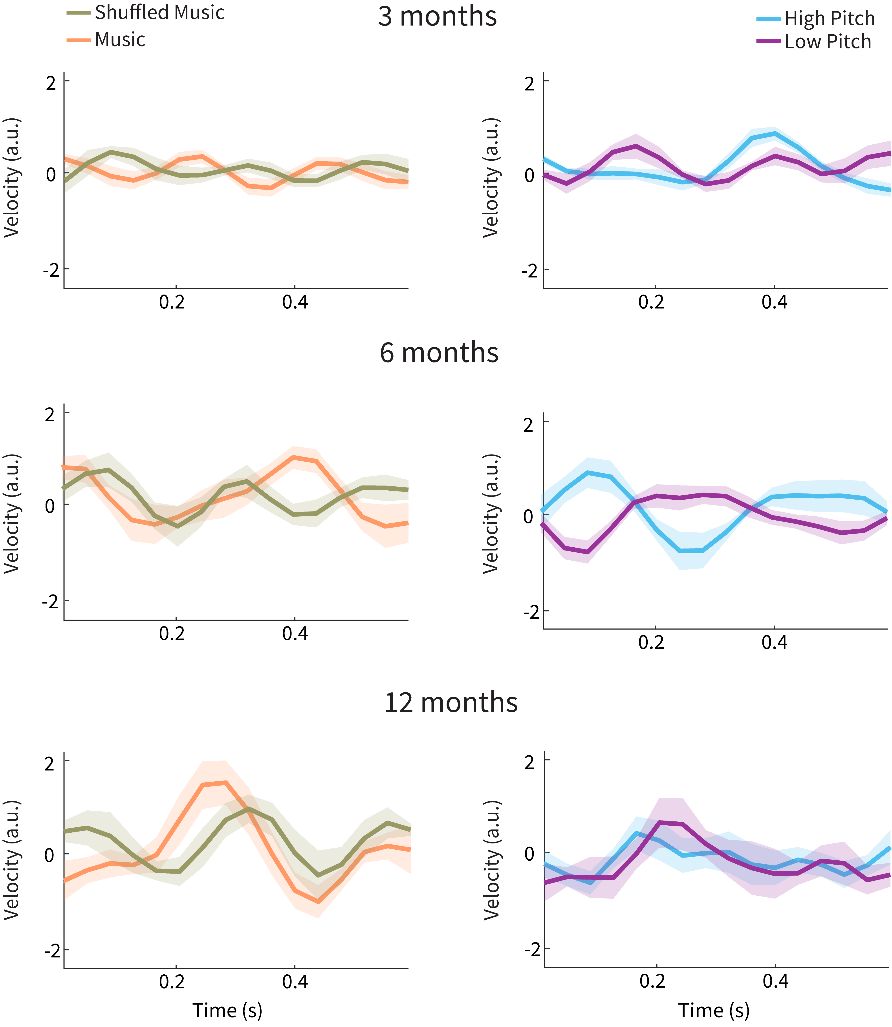

***Figure S3****.* *The figure displays the z-scored velocity (a.u., y-axis) of infants’ movement responses over time (in seconds, x-axis) following peaks in the amplitude envelope of the stimuli (at time = 0) across different conditions. These peaks represent moments of increased acoustic intensity in the music, allowing visualization of how infants at three developmental stages (3 months, top row; 6 months, middle row; 12 months, bottom row) synchronized their movements with dynamic changes in the sound. Each subplot compares different conditions, with left plots comparing music (orange) to shuffled music (khaki) and right plots comparing high-pitch (blue) with low-pitch music (purple) conditions. None of the age groups show significant differences in velocities following peaks in the amplitude envelope of the auditory stimuli across conditions. Specifically, the results suggest that changes in sound intensity seem to evoke movement responses that vary in latency and might be inconsistent across trials (with the only exception of the 12-month-old infants, who appear to show a peak around 200 ms across several conditions). In this context, such varying responses are better captured by Granger Causality analysis. Perhaps because such predictive approaches are more powerful in picking up signals that might be either sporadic or variable in amplitude and/or sign across trials (Shojaie & Fox, 2022).*

**Movement – Periodicity analyses**

Despite infants showing no consistent phase-locked movement responses, we tested further whether these responses were periodic and, if so, whether such periodicity differed across conditions. To do so, we used an auto-correlation approach described in a previous study reporting periodic movements in infants exposed to music (Zentner & Eerola, 2010). The analysis builds upon the distribution of the highest coefficients yielded by autocorrelation analyses (*Fig. S3*). We first examined whether the density distribution of movements was uniform (indicating non-periodic movements) or non-uniform (indicating periodic movements) across PMs for each age group. To do so, we performed a Kolmogorov-Smirnov test, which revealed a highly significant result (*KS*=0.838-0.869, *p*<.001), indicating that the movements were periodic across all PMs and age groups. Next, we extracted the density value associated with the beat-related lag (444 ms) and compared these values across conditions averaged over all PMs per age group. Results indicated no significant differences in periodicity between music and shuffled music (*ps*>.120). All infants, therefore, demonstrated periodic motor responses to auditory stimuli, regardless of whether they listened to music or shuffled music. Additional analyses comparing the pitch conditions revealed no significant differences either (*ps*>.095).

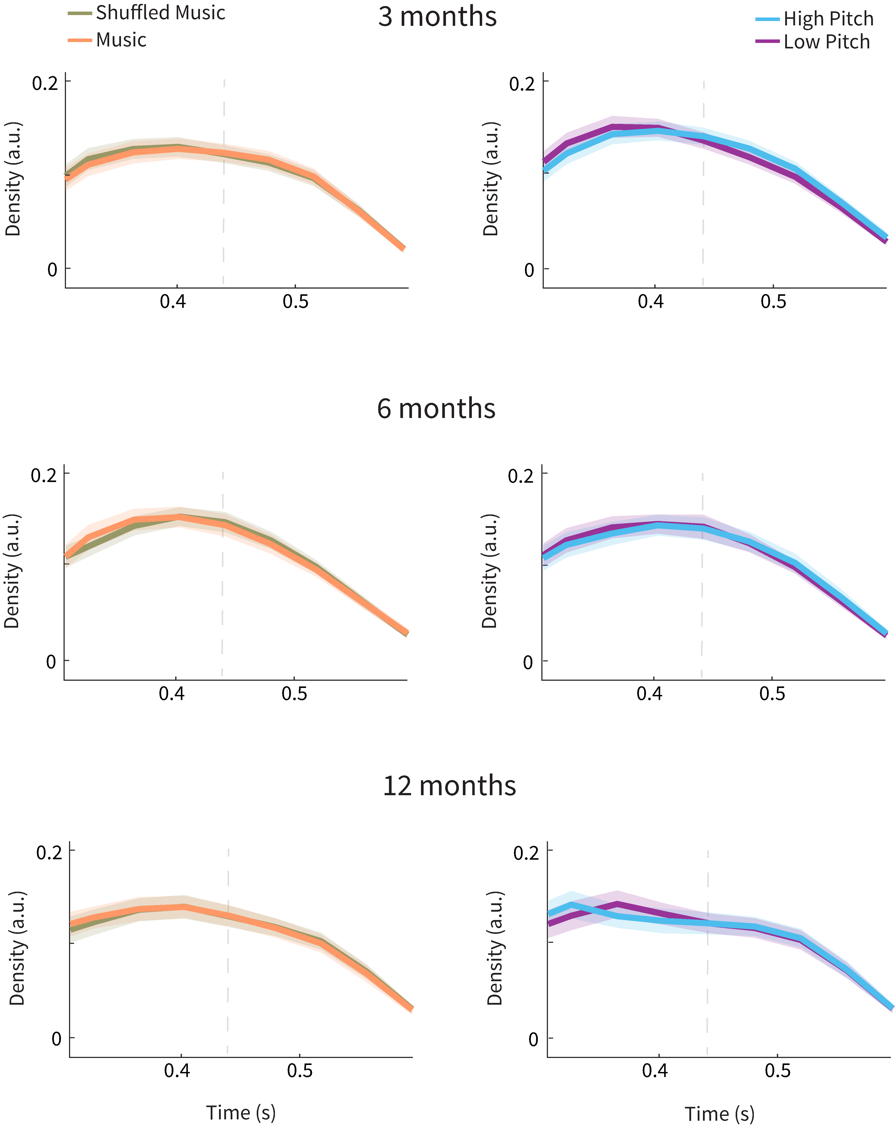

***Figure S4***. *The graphs depict the density (y Axis) function of infants’ periodic movement to music (orange) vs shuffles music (khaki) as well as high-pitch (blue) vs low-pitch music (purple) averaged across all PMs at three developmental stages: 3 months (top row), 6 months (middle row) and 12 months (bottom row). The lag in seconds is depicted on the x Axis and the dotted vertical line indicates the musical beat (444 ms lag). Infants’ movements tended to be rhythmic, however, not differently across conditions (i.e., the contrasts of density values across conditions were not significant).*

**Links between neural and movement responses**

Additional exploratory analyses examined correlations between ERP amplitudes and movement measures using participant-averaged data. Mean ERP amplitudes of the P1 component time window, identified from cluster-based analyses, served as neural measures. Movement measures included (1) total movement quantity (mean velocity across the trial) and (2) Granger causality F-values indicating music-to-movement coupling strength. Analyses contrasted music versus shuffled music conditions and high-pitch versus low-pitch conditions. Linear mixed-effects models were fitted with ERP amplitude as the response variable, either movement quantity or Granger causality F-values as fixed effects, and infants as random intercepts. Results revealed no significant correlations between ERP amplitude and movement quantity, irrespective of conditions (*p*>.124), and neither when comparing music versus shuffled music (*p*>0.111) or high versus low pitch (*p*>0.071). Similarly, no significant correlations were found between ERP amplitude and Granger causality F-values, irrespective of conditions (*p*>.164) and in either contrast (music vs. shuffled music: *p*>0.494; high vs. low pitch: *p*>0.175). These findings might suggest that neural sensitivity to musical structure and motor responsiveness develop independently within the first year of life, consistent with developmental theories positing perceptual sensitivity precedes motor coordination.
